# Motile curved bacteria are Pareto-optimal

**DOI:** 10.1101/441139

**Authors:** Rudi Schuech, Tatjana Hoehfurtner, David Smith, Stuart Humphries

## Abstract

Curved-rods are a ubiquitous bacterial phenotype, but the fundamental question of why they are shaped this way remains unanswered. Through *in silico* experiments, we assessed freely swimming straight- and curved-rod bacteria of a wide diversity of equal-volume shapes parameterized by elongation and curvature, and predicted their performances in tasks likely to strongly influence overall fitness. Performance tradeoffs between these tasks lead to a variety of shapes that are Pareto-optimal, including coccoids, all straight rods, and a range of curvatures. Comparison with an extensive morphological survey of motile curved-rod bacteria indicates that the vast majority of species fall within the Pareto-optimal region of morphospace. This result is consistent with evolutionary tradeoffs between just three tasks: efficient swimming, chemotaxis, and low cell construction cost. We thus reveal the underlying selective pressures driving morphological diversity in a wide-spread component of microbial ecosystems.

**Significance Statement:** Bacteria exhibit a bewildering diversity of morphologies but despite their impact on nearly all aspects of life, they are frequently classified into a few general categories, usually just ‘spheres’ and ‘rods’. Curved-rod bacteria are one simple variation and are widespread, particularly in the ocean. However, why so many species have evolved this shape is unknown. We show that curvature can increase swimming efficiency, revealing a widely-applicable selective advantage. Furthermore, we show that the distribution of cell lengths and curvatures observed across bacteria in nature are predicted by evolutionary tradeoffs between three tasks influenced by shape: efficient swimming, the ability to detect chemical gradients, and reduced cost of cell construction. We therefore reveal shape as an important component of microbial fitness.

## Main Text

In stark contrast to the macroscopic realm of multicellular plants and animals, in the microscopic world it remains a challenge to attribute function to any of the considerable morphological diversity that exists (1, 2). Even superficially simple traits such as the curvature of many rod-shaped bacteria have eluded evolutionary explanations, despite the ubiquity of such morphological variations. For example, while coccoids and “rods” comprise the vast majority of bacteria in the ocean, curved rods account for up to a quarter of these (3), sometimes outnumbering straight rods (4). Common in most other habitats (5–9), curved bacteria have invested heavily in complex genetic machinery to generate and maintain their shapes (10), but we have almost no understanding of the benefits their curvature confers (1, 11). While the biochemical and genetic mechanisms of cell growth and the maintenance of shape have received considerable attention (12–15), the costs or benefits of a particular shape have received almost none (but see (16, 17)). Understanding the links between form and function is a cornerstone of modern biology, yet no studies have addressed why curvature has evolved in many bacterial clades (5–9).

The microscale physics relevant to bacteria makes addressing the physical consequences of their shapes particularly non-intuitive (18). One hypothesis is that at microscopic scales, shape generally has no bearing on fitness (19). However, while there appears to be limited support for this ‘neutral morphology’ theory (20), several lines of empirical evidence suggest that, just as for larger organisms, selective pressures influence microbial cell shape. Bacterial cell shape is 1) heritable, with some morphologies evolving independently many times; 2) has high diversity, yet is typically uniform (excluding pleomorphism) within species; and 3) is known to be actively modified in response to environmental changes (1).

Free-swimming cells are ubiquitous, comprise a large fraction (up to 70%) of the bacteria in the oligotrophic marine environment (21, 22), and are part of the life cycle of many biofilm-forming species. However, the streamlining principles that explain the shapes of many swimming animals work very differently at the microscale (23). The most efficient shape for an ellipsoidal swimmer at this scale is similar to a rugby ball (18, 24), but most rod-shaped bacteria are more elongated than this optimum (25). Dusenbery (25) suggests that motile bacteria are under selective pressure to elongate without bound in order to improve chemotactic ability, since such cells are much more resistant to random course changes imposed by Brownian motion. Hence, the diversity of elongated shapes in bacteria may well be due to evolutionary trade-offs between a number of tasks (26), such as chemotaxis and swimming efficiency. Might bacterial curvature have evolved due to similar tradeoffs?

Here, we address a broad bacterial morphospace of all free-swimming curved rods, straight rods, and spheres. First, we survey the literature in search of microscopy data on curved bacteria, inserting them into a two-dimensional parameter space of elongation and curvature to quantify their morphological diversity. In a major effort involving tens of thousands of numerical simulations, we then quantify performance throughout this morphospace in several physiologically relevant tasks likely to contribute to overall evolutionary fitness, revealing the selective advantages (and disadvantages) of curvature and elongation. Finally, from these disparate performance landscapes*, we use the concept of Pareto optimality (26) to analyze the diversity of observed rod morphologies through the lens of evolutionary trade-offs between tasks. While previous work (26) has approached this type of problem by leveraging abundant data on the ecology of macroscopic organisms (e.g. birds, bats, ants), such data is sparse or non-existent at the scales relevant to bacteria. Despite this major obstacle, our direct computations of Pareto optimality allow us to identify which tasks likely constrain the evolution of shape within this morphologically diverse polyphyletic group. We thus link form and function at the previously intractable microscale and pave the way toward a broader understanding of bacterial diversity.

## Survey of curved bacteria shapes

We surveyed the literature (5–9, 28) for micrographs of motile, flagellated curved rod bacteria (*Methods*, SI); since a wide range of motile straight rods (from coccoids to filaments) has previously been reported (25), we implicitly include these in our subsequent calculations. Individual cells were segmented using MicrobeJ (29) and fitted to a simple geometric model parameterized by elongation (*ℒ*) and centerline “curvature” (*𝒦*)^†^, both dimensionless and thus independent of size (Fig. 1 inset, *Methods*). The survey (Fig. 1, Table S3) highlights the diversity of shapes within our curved rod parameter space: short bean-, comma-, and long hotdog-like shapes can all be found in nature, with a few exceptionally elongated species beyond *ℒ* = 10 (Fig. S 1). Of the 11 phyla, 98 genera and 205 species, notable members of the dataset include *Bdellovibrio* spp. (which feeds on other bacteria, including human and animal pathogens), and *Vibrio cholerae* and *V. vulnificus*, both capable of causing serious illness in humans. The median curved rod (*ℒ* = 3.7, *𝒦* = 0.14) corresponds to a rod with limited curvature (similar to *Vibrio ruber*), and 50% of all curved species fall within *ℒ* = 2.3 to 4.9 and *𝒦* = 0.05 to 0.25. There is a factor of ten variation in equivalent spherical diameter (ESD), but with no discernible correlation between size and shape (*ℒ* vs ESD, *F_1,218_* = 0.303, *p* = 0.582; *𝒦* vs ESD, *F_1,218_* = 1.089, *p* = 0.298), we do not consider size further (SI). There is, however, a weak but decreasing trend in curvature versus elongation (*ℒ* vs *𝒦*, *F_1,221_* = 6.360, *p* = 0.03, *r^2^* = 0.012). Although toroidal bacterial morphologies do exist‡, there is a conspicuous lack of motile species that are more than semi-circular (Fig. 1).

**Fig. 1.**
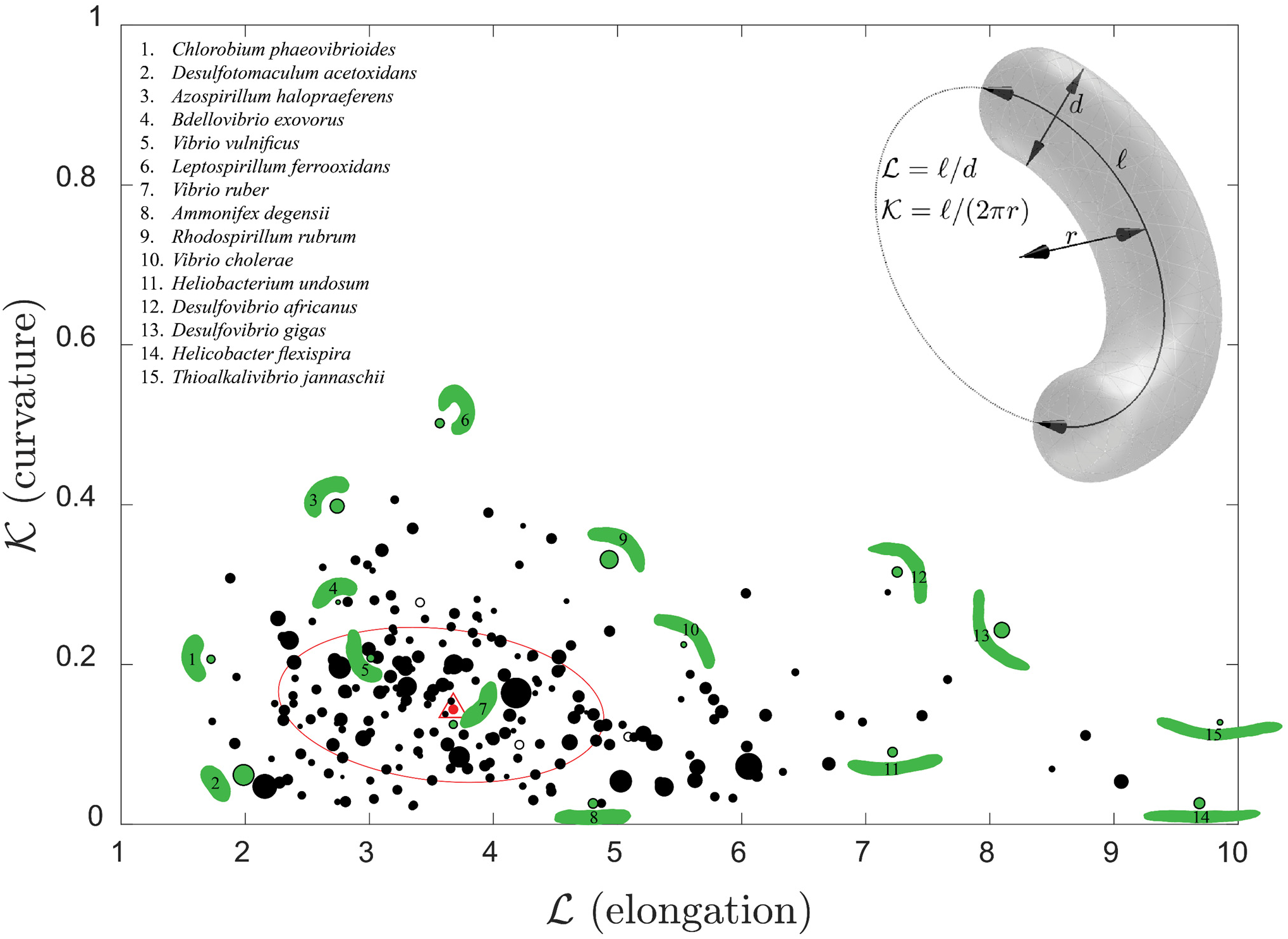
Survey of extant curved rod morphologies, with geometric parameters and definitions of dimensionless shape parameters *ℒ* and *𝒦* (inset). Circles represent species median shapes based on segmented images, with circle radius proportional to equivalent spherical diameter when size data available (filled) or overall median size when not (open). Smallest ellipse containing 50% of all species is shown in red, and overall median species shape (and size) is depicted by a red filled circle, highlighted by a triangle. Silhouetted individuals (green), rescaled to constant volume, are shown for selected species (green filled circles).

## Swimming Efficiency

Given that curved bacteria fill a particular region of the theoretical morphospace (Fig. 1), it is natural to ask why bacteria might be curved at all, why some are as curved as they are, and why not all conceivable curved rod morphologies are found in nature. We first hypothesized that cell curvature might enhance Swimming Efficiency **ψ**_*swim*_, defined as the ratio of power required to translate an equal-volume sphere to the mechanical power dissipated by the flagellar motor (*Methods*, SI). This hypothesis is in line with the results of Phan Thien et al. (34), who showed that a slightly flattened triaxial ellipsoid is marginally more efficient than the best rotationally symmetric spheroid, as well as Liu et al. (35), who approximated *Caulobacter crescentus* as a straight rod with a tilted off-axis flagellum and found increased efficiency compared to the aligned case. The reason for this is a peculiarity of propulsion by a rotating flagellum - there is a tradeoff between minimizing the translational resistance of the body and maximizing its rotational resistance, to reduce the power typically wasted by body counter-rotation. Applied to curved rods, this idea suggests that an optimal curvature should exist, since curvature should increase both translational and rotational resistance.

Using a regularized Stokeslet Boundary Element Method (*Methods*, SI), we simulated the detailed kinematics of freely-swimming curved rods (Fig. 2B, Movie S1), generating a “performance landscape” for Swimming Efficiency (Fig. 2A) over a broad theoretical morphospace spanning nearly all our observed body shapes as well as spheres, straight rods, and rings (Fig. 2A, Fig. S 12). All shapes were equal-volume (1 μm^3^, similar to *E. coli*) to focus on variation in shape, not size.

**Fig. 2.**
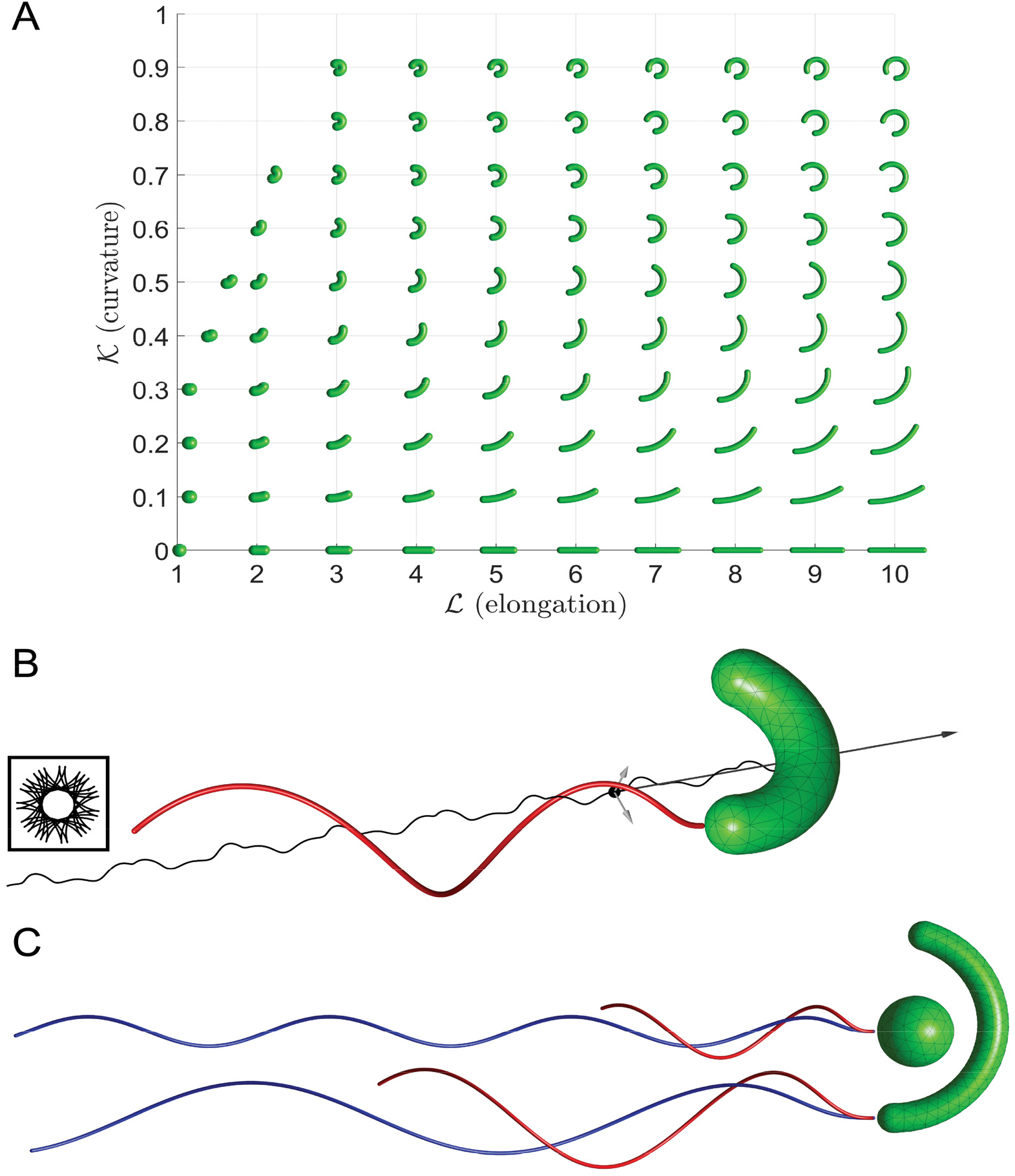
A) Simulated two-dimensional morphospace of equal-volume body shapes. Empty region at upper left consists of nonphysical self-intersecting shapes. B) Example body + flagellum with simulated swimming trajectory traced by the body midpoint, which appears to lack any symmetry viewed off-axis but reveals long-range rotational symmetry viewed axially (inset). The random Brownian rotation that would be superimposed onto this swimming trajectory can be quantified by three principal rotational diffusivities, depicted as vectors originating from the center of diffusion (black circle) **(36)**. The largest of these (dark gray) corresponds to rotations around the flagellar axis, but it is the other two (light gray) that determine how long the cell can maintain its course. C) Comparison of optimal flagellar shapes for a sphere (*ℒ* = 1, *𝒦* = 0) and highly elongated curved rod (*ℒ* = 10, *𝒦* = 0.5), for Swimming Efficiency (red) and Chemotactic SNR (blue). Example 2^nd^ order triangular surface meshes are shown in B and C; the flagella were similarly fully meshed (SI).

Surprisingly, we found that over the entire morphospace, the globally optimal Swimming Efficiency is achieved not by a curved rod but a straight, slightly elongated rod comparable to a medicine capsule (*ℒ* = 1.46, *𝒦* = 0; Fig. 3A). However, this shape has a Swimming Efficiency advantage of just 2% over a spherical body, in general agreement with previous studies of ellipsoidal cells (18, 24, 34) (Fig. S 16), and is only 0.3% better than the semicircular curved rod located at a local maximum in the performance landscape (*ℒ* = 4.0, *𝒦* = 0.70). In fact, a diverse range of shapes exhibit essentially the same efficiency, with only long straight rods having substantially reduced performance compared to a sphere (Movie S1).

Intriguingly, the optimal curvature *𝒦* for a given elongation *ℒ* is approximately constant (~0.65) beyond *ℒ* = 4, with efficiencies within a few percent of that of a sphere up to *ℒ* = 10. However, even modest curvature (e.g. *𝒦* = 0.2) allows a highly elongated straight rod (*ℒ* = 10) to mitigate its efficiency penalty by over 20%. Thus, if bacteria were to elongate for reasons other than efficient swimming, they could avoid this penalty by being curved. While Swimming Efficiency (or equivalently, speed – see SI) alone cannot explain why elongated, curved bacteria have evolved, these data suggest efficiency might nonetheless apply selective pressure if it were to influence some other ‘higher-level’ task favoring elongated cells.

**Fig. 3.**
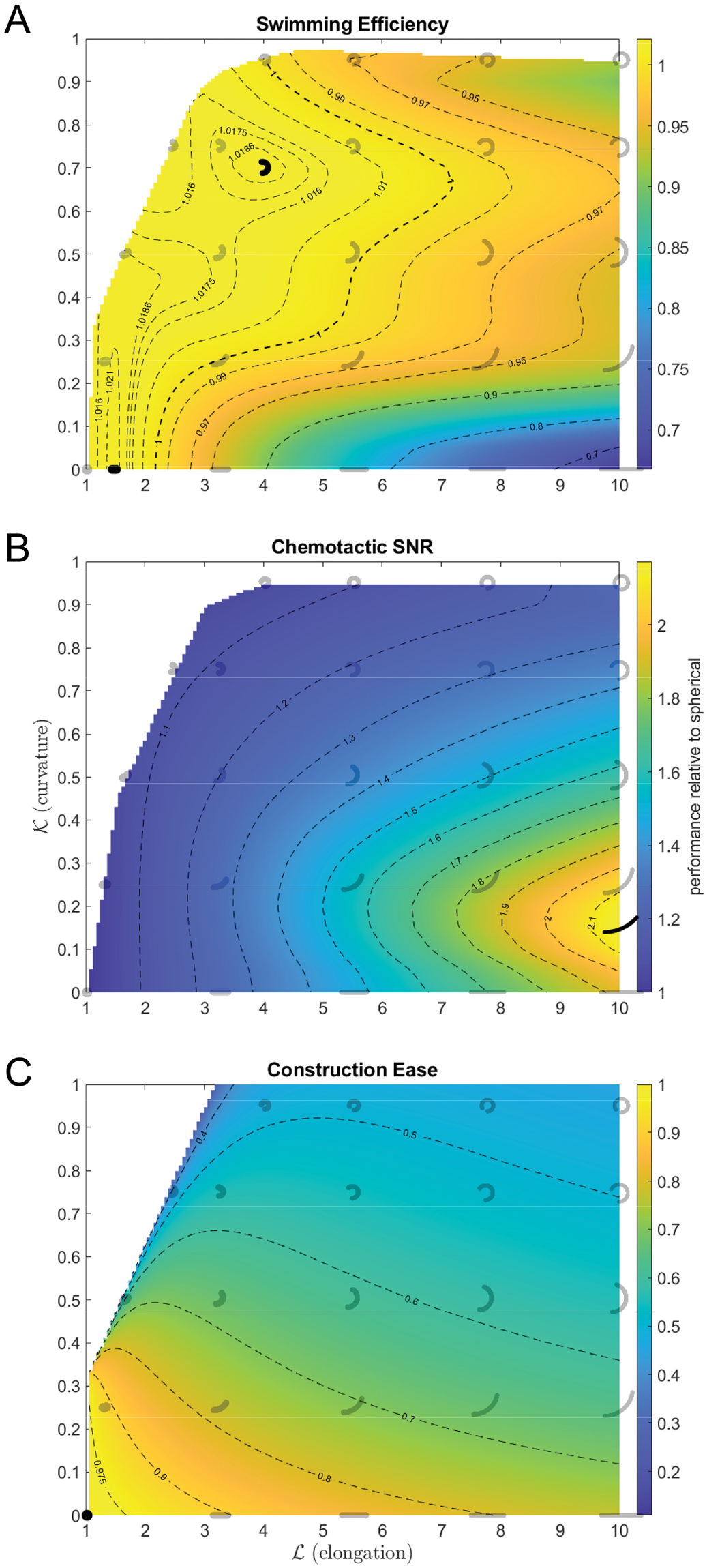
Performance landscapes for putative tasks critical to curved rod bacteria: Swimming Efficiency 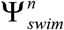 (A), Chemotactic SNR 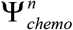 (B), and body shape Construction Ease 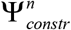 (C). In each case, performance (pseudocolor and contours) is normalized relative to that of a spherical body (*ℒ* = 1, *𝒦* = 0) and is thus dimensionless. In addition to selected shapes for reference (gray), performance maxima within our morphospace are shown (black); in panel A, these include both the global and a local maximum (see text).

## Chemotactic Signal/Noise Ratio

It is commonly assumed that motile bacteria swim to enable chemotaxis (37, 38) – the directed movement of organisms toward high concentrations of favorable compounds (e.g. nutrients) or away from unfavorable ones. To perform chemotaxis, bacteria must reliably sample chemical concentration at different points in space to determine whether the stimulus gradient is increasing or decreasing as they swim. For ellipsoidal cells, Dusenbery (25) concluded that many bacteria have likely evolved elongated shapes due to the benefits to chemotactic ability. The advantage of elongation here derives from larger resistance to random Brownian rotation – the longer a bacterium can maintain its orientation, the longer it can trust its concentration gradient estimates before Brownian motion randomizes its direction of travel. Following Dusenbery’s information theory approach, we quantified how the reliability, or signal to noise ratio (SNR), for these gradient estimates varies with elongation *ℒ* and curvature *𝒦* of rod-shaped bacteria.

We note that overall chemotactic ability depends on other factors in addition to SNR. For instance, active reorientation is important (39, 40), and is observed as “tumbles” in *E. coli* and “flicks” in *Vibrio* spp. (41). Hence, we also considered performance in a task that approximates the effect of shape on the ability to reorient – Tumbling Ease (SI). In addition, since a primary purpose of chemotaxis is to maximize a cell’s ability to obtain nutrients, we investigated the effect of shape on Nutrient Uptake via diffusion to the cell surface (SI). However, (discussed later) we found that in contrast to chemotactic SNR, these two related tasks were unnecessary to explain the diversity of rod morphologies.

For a bacterium comparing concentration samples over time, Chemotactic SNR is proportional to 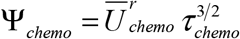, where *τ_chemo_* is the sampling time or timescale for loss of orientation and 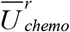 is the effective swimming speed (*Methods*, SI) (18). Chemotactic SNR will thus benefit from both increased resistance to Brownian rotation (which increases *τ_chemo_*) as well as increased Swimming Efficiency (which increases 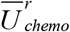), so one might hypothesize that the optimal shape for Chemotactic SNR would be highly elongated as well as curved. We numerically calculated anisotropic rotational diffusivities (Fig. 2B, SI) to quantify how *τ_chemo_* varies with shape, and calculated 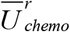 via simulations similar to those employed for Swimming Efficiency (*Methods*, SI).

We find that Chemotactic SNR also increases without bound with elongation *ℒ* (albeit slower than previous predictions, Fig. S 7) due to a faster increase in *τ_chemo_* (Fig. S 4) than decrease in 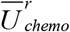 (Fig. S 6), so that there is no finite optimal cell shape for this task (Fig. 3B). Crucially, however, there is a non-zero optimal curvature *𝒦* for any *ℒ* due to another tradeoff between *τ_chemo_* (which favors straight rods, Fig. S 4) and 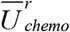 (which tends to favor substantial curvature, Fig. S 6). Thus, within our simulated morphospace, the best shape for Chemotactic SNR is a slender, slightly curved rod (*ℒ* = 10, *𝒦* = 0.15; Fig. 3B). Selection for Chemotactic SNR can therefore qualitatively explain why most rod-shaped bacteria are more elongated than the pill-shaped Swimming Efficiency optimum (18) as well as being curved. However, bacteria obviously cannot elongate without bound as selection solely for SNR would suggest. In addition, although we focused on surveying curved rods, many straight rods along a continuum of elongation also occur (25) and most of these presumably should not exist if the only relevant selective pressures were Swimming Efficiency and Chemotactic SNR.

## Construction Ease

One reason for the existence of straight rods might be that they are simply easier to construct than curved cells. In fact, the least costly body shape is likely to be spherical (10, 42), with specific structural proteins required to maintain both rod (43) and vibrioid or helical (44) cell shapes. It therefore seems reasonable to assume the existence of a selection pressure that penalizes morphologies with sharply curving surfaces (18, 45). Building on Dusenbery’s assumption of a minimum feasible radius of curvature (18), and in the absence of a general theory for predicting the cost of constructing an arbitrarily shaped cell wall, we considered a number of geometrical cost functions (SI) based on common metrics of the principal curvatures *k*_1_, *k*_2_. The simplest cost function consistent with our observed bacterial shapes (SI) is based on total absolute Gaussian curvature weighted by the area *A* of the body surface *S*, which we inverted to yield 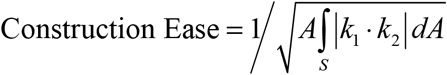; see *Methods* for more details.

As intuitively required, the maximum Construction Ease is achieved by a coccoid cell (*ℒ* = 1, *𝒦* = 0; Fig. 3C), with both elongation *ℒ* and curvature *𝒦* generally increasing costs§. For almost all *ℒ*, the dependence on *𝒦* is stronger than on *ℒ* so that in this putative task, curvature is costlier than elongation. While total absolute Gaussian curvature is constant for all straight rods, the area penalization ensures that elongation when *𝒦* = 0 remains costly. Thus, Construction Ease should exert selective pressure on bacteria to minimize both elongation and curvature over evolutionary time.

## Performance tradeoffs and Pareto optimality

Our results quantify the relative advantages and disadvantages of bacterial curvature and elongation. Relative to straight rods of equal elongation *ℒ* within our morphospace, curved rods of median curvature *𝒦* = 0.14 exhibit up to a 27% improvement in Swimming Efficiency and up to 12% improvement in Chemotactic SNR, but a 15 - 40% disadvantage in Construction Ease (Fig. S 17). In reality, these performance tradeoffs occur in two dimensions: a curved rod of *ℒ* = 5, *𝒦* = 0.2 will simultaneously experience pressure to straighten and shorten (to increase Construction Ease, Fig. 3C), to elongate (to increase Chemotactic SNR, Fig. 3B), and to become *more* curved (to increase Swimming Efficiency most rapidly, Fig. 3A).

Faced with such complex trade-offs between tasks, how can one approach the question of optimal shapes? The overall fitness of an organism can be assumed to be an increasing function of the performances of all tasks, but this fitness function is rarely known. Shoval et al. (26) discuss an elegant solution to this multi-objective optimization problem: Pareto optimality theory. This concept, borrowed from economics and engineering, provides a set of solutions that are the best trade-offs between individual tasks; within this optimal set, it is impossible to improve at one task without sacrificing performance at another. In the context of morphology, Pareto-optimal shapes can be contrasted with sub-optimal shapes that can be outperformed in all tasks simultaneously, and which are not expected to occur naturally. The shape of any given Pareto-optimal species depends on the relative contributions of each task to overall fitness in its ecological niche, i.e., the particular fitness function of that niche (26).

While Pareto tradeoffs between a small number of critically important tasks often explain the majority of phenotypes seen in nature (26), we did not know *a priori* what these tasks might be for motile rod-shaped bacteria. Hence, using a simple brute-force approach, we directly computed the Pareto-optimal regions of morphospace from the performance landscapes of many possible combinations (sets of 3 – 5) of our putative tasks (i.e., Swimming Efficiency, Chemotactic SNR, Construction Ease, Tumbling Ease, and Nutrient Uptake, see *Methods*, SI). We inferred the most likely set of crucial tasks by identifying the best goodness-of-fit (GoF) metric between each theoretically optimal set of shapes and the morphospace region bounded by observed species (*Methods*, SI).

The best GoF** corresponds to a three-way tradeoff between Swimming Efficiency, Chemotactic SNR, and Construction Ease (SI), indicating that these three tasks have been critically important in shaping the evolution of motile curved rods. The resulting Pareto-optimal region (Fig. 4, SI) reproduces the decreasing trend in maximum curvature *𝒦* of observed morphologies and encompasses the diversity of species shapes, with most bacteria apparently employing a generalist strategy but some species specializing in just one or two tasks. In particular, the boundaries (i.e., fronts) of the main Pareto-optimal region consist of bacteria engaged in tradeoffs between only two tasks: Construction Ease versus Chemotactic SNR for most straight rods along the elongation (*ℒ*) axis (e.g. *Ammonifex degensii*), and Swimming Efficiency versus Chemotactic SNR along most of the upper boundary (e.g. *Rhodospirillum rubrum*). There are only two theoretically suboptimal outliers: *Desulfovibrio africanus* (*ℒ* = 7.3, *𝒦* = 0.32), and *Leptospirillum ferrooxidans* (*ℒ* = 3.6, *𝒦* = 0.50), which often has an extreme C-shape and erodes ferrite surfaces (46), possibly facilitating high contact area with its substrate.

**Fig. 4.**
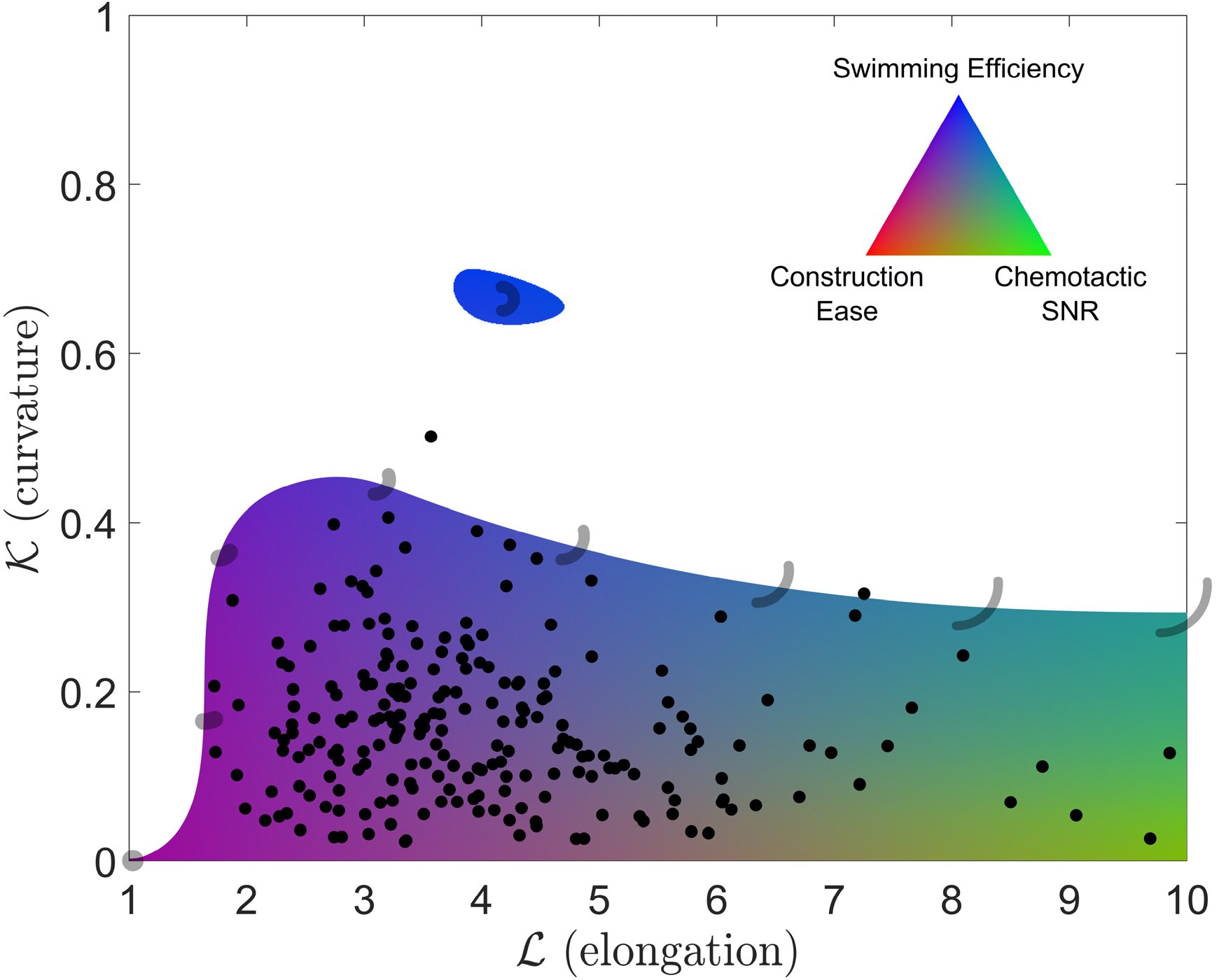
Pareto-optimal curved rod morphologies. Colored regions represent the set of shapes that are Pareto-optimal resulting from tradeoffs between Swimming Efficiency, Chemotactic SNR, and Construction Ease, white region represents sub-optimal shapes, and dots represent observed species medians. Selected simulated morphologies are plotted along the main optimal/sub-optimal Pareto front as well as at the centroid of the disconnected Pareto-optimal “island.” RGB color values were assigned by normalizing performances in each task within the optimal region between 0 – 1. Color thus signifies relative, not absolute, tradeoffs between tasks as indicated by the color triangle (inset). Not all colors in the triangle are realized because not all possible tradeoffs are realized (e.g. the shapes that excel at Construction Ease also excel at Swimming Efficiency).

Crucially, the Pareto-optimal region excludes almost all shapes that are not seen in nature, e.g. short, sharply curved cells and motile ring-shaped cells. However, we note the conspicuous presence of a small “evolutionary island” of semicircular rods due to the nearby relative maximum in the Swimming Efficiency performance landscape (Fig. 3A). While these shapes may be theoretically optimal, their absence in nature (at least from our survey) could simply be a consequence of the vast “sea” of suboptimality isolating them from observed morphologies. We also predict, but do not observe, the existence of highly elongated, moderately curved rods (e.g. *ℒ* = 10, *𝒦* = 0.3). However, in reality all curved rod bacteria may have a three-dimensional helical shape that is difficult to discern from typical microscopy images (10, 47). We suspect that the likelihood of cell curvature appearing clearly helical and not planar increases with *ℒ*, so that such long, curved rods do exist in a sense, but are helically shaped and beyond the scope of this study.

## Discussion

We modeled performance in several putative tasks likely to be important to motile bacteria, in some cases with little *a priori* knowledge of which morphologies might optimize them. Our detailed numerical simulations reveal performance landscapes with complex topology (i.e., non-elliptical contours (Fig. 3A,C), local maxima (Fig. 3A), non-finite archetypes (Fig. 3B), which combine to yield a Pareto-optimal region that is not a simple triangle (Fig. 4) as would be expected under simplifying assumptions (26). By considering many combinations of the putative tasks in our analysis of Pareto-optimality, we gain confidence** in our conclusion that the tasks constraining evolution of curved morphologies are Swimming Efficiency, Chemotactic SNR, and Construction Ease. While we cannot rule out the importance of additional tasks such as navigation through viscous gels (17), avoidance of predation (48, 49), attachment or detachment from surfaces (16, 50), and activities related to life in biofilms (51), our proposed set of three tasks is minimally complex in the sense that this is the minimum number required to explain the diversity of observed morphologies (SI).

The inclusion of Swimming Efficiency in the small set of critically important tasks shaping the evolution of curved bacteria might seem surprising given the ranges of variation seen in the performance landscapes: Construction Ease and Chemotactic SNR display relatively strong effects of shape, with factors of 3 and 2.2 variation, respectively, over the Pareto-optimal region of morphospace (Fig. 3B, C). Although Swimming Efficiency exhibits a factor of 1.5 variation overall, there is far less variability (i.e., 10%) over the majority of considered shapes, particularly those that determine the upper front of the main Pareto-optimal region (Fig. 3A, Fig. 4). This observation highlights a core difficulty with the evolutionary multi-objective optimization problem – the relative contributions of each task to overall fitness are unknown, so that slight changes in traits such as Swimming Efficiency may have a disproportionately large impact on fitness (52). Furthermore, natural selection operates primarily on tiny variations in fitness that are difficult or impossible to observe experimentally (53). Lastly, Swimming Efficiency is a fundamental motility parameter and is likely to contribute to many other higher-level tasks, so its effect on overall fitness is probably compounded. For instance, not only is Chemotactic SNR enhanced by curvature due to its dependence on Swimming Efficiency, but overall chemotactic ability will depend on how quickly the cell can swim toward better conditions, in addition to the reliability of its gradient measurements. Experiments with marine bacteria, which tend to be fast-swimming but live in a nutrient-poor environment (41), have confirmed a strong relationship between swimming speed and overall chemotactic ability (54).

The diversity of bacterial morphologies found in nature is reminiscent of the perhaps even larger diversity in marine phytoplankton, which prompted the “the paradox of the plankton” posed by Hutchinson (55). The paradox asked how there can be such a wide range of phytoplankton species, all competing for the same resources in the same habitat, despite the competitive exclusion principle predicting that one species should eventually out-compete the rest. Many resolutions of the paradox note that the open ocean is not a homogeneous habitat as originally thought, but quite spatially and temporally complex at small scales, so that species actually occupy different ecological niches (56). Similarly, while the diversity of bacterial morphologies coexisting in environments such as biofilms (51, 57) or the human gut (58) might seem paradoxical, this level of habitat classification is far too coarse to be relevant to bacteria. In contrast to macroscopic organisms (59), our understanding of the niches occupied by bacteria is extremely limited. Nonetheless, our results provide testable predictions for which tasks, and perhaps micro-niches, different rod-shaped morphologies are specialized. For example, highly elongated species should display very effective chemotaxis and therefore might be adapted to habitats with shallow nutrient gradients. Species with high curvature, being efficient swimmers, might tend to live in oligotrophic or highly viscous environments. The low cost of construction we predict for nearly spherical bacteria is difficult to observe but one might speculate a correlation between ease of construction and growth rate. While controlling for the many confounding factors that vary across species is challenging, we anticipate that advances in experimental (60) and phylogenetic methods (61) will enable direct tests of these predictions.

## Methods

### Survey of observed shapes

We examined all images in Bergey’s Manual ^®^ of Systematic Bacteriology (5–9) and The Prokaryotes (28) for curved rods. We additionally filtered an in-house manually collected dataset of bacterial morphologies compiled according to the methods of (62) for all species that were reported as “curved,” “slightly curved,” “bent,” “kidney-shaped,” “crescent-shaped,” “comma-shaped,” “crooked,” “C-shaped,” or “boat-shaped.” We then searched for these species (via genus) on http://www.bacterio.net/ and scanned the publications listed there for images of cells; if this failed, we tried a Google Scholar search for publications on the species. We restricted our searches to bacteria identified to species level, known or likely to exhibit swimming motility (i.e. flagellated, or swimming motility demonstrated for all other members of genus), and to images that contained at least some perceptibly curved individuals; see SI for full details of the methodology. In all, our dataset consists of 4903 individual bacteria across 363 images (Table S2).

Our morphology measures (elongation *ℒ* and curvature *𝒦*) were determined after semi-automatically segmenting the cells using the “Rod-Shaped” shape type in MicrobeJ (29) (SI). Each segmented image was manually inspected to ensure accuracy of all measurements.

We computed unweighted species medians of *ℒ*, *𝒦*, and equivalent spherical diameter by first taking medians within each image, aggregating multiple images into one dataset for this purpose when they were stated to represent identical conditions (e.g. strain, substrate, nutrient conditions, etc.). We then calculated the medians of these values across all images found for each species, regardless of strain (Table S3). Using means instead of medians yielded similar results and did not affect our conclusions.

To calculate Goodness of Fit (GoF) between the theoretically Pareto-optimal regions and observed bacterial shapes, we computed a polygonal boundary similar to a convex hull around the latter data points; see SI for more details.

### Model geometry

The bacterial cell body was modeled as a curved rod of circular cross section with hemispherical caps on both ends. It has been suggested that all curved rod bacteria are actually short sections of a three dimensional helix (47) but for simplicity we assume that the cell centerline always lies in a plane. All cells had a constant, arbitrary (see SI) volume of 1 μm^3^, so each body shape was determined by two dimensionless aspect ratios: *ℒ* = ℓ/*d*, which describes elongation, and *𝒦* = ℓ (2*πr*), which describes the degree of centerline curvature of the curved rod shape, where ℓ is the total arclength of the body centerline (from pole to pole), *d* is the body diameter, and *r* is the radius of curvature of the body centerline (Fig. 1 inset).

The flagellum was modeled as a modified right-handed helix of finite circular cross section (radius 0.03 μm) parameterized by amplitude *a*, wavelength λ, and number of wavelengths *n_λ_*, as in Shum et al. (24). Full details are given in SI.

### Numerical solution for swimming kinematics and rotational diffusion

A regularized Stokeslet Boundary Element Method was used to numerically solve for the inertia-less kinematics of freely swimming cells propelled by a rotating flagellum in water, i.e. their effective swimming speed *Ū;_swim_* and average power dissipation by the flagellar motor *P̄^M^*. This type of numerical method has been used extensively to model swimming microorganisms; our approach generally follows that of Smith (63) with several performance improvements detailed in SI. We also used this code to quantify the anisotropic Brownian diffusion (36) of curved rod bacteria, accounting for the stabilizing effect of the flagellum (64). Specifically, we calculated the timescale for loss of orientation *τ*, which depends on the rotational diffusion coefficients for rotations around the two principal axes normal to the swimming direction (18) (Fig. 2B, SI).

### Putative tasks

#### Swimming Efficiency

Swimming Efficiency was defined as 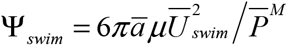, where *â* is equivalent cell radius (0.62 μm for all cells here) and *μ* is dynamic viscosity of water (10^-9^ kg s^-1^ μm^-1^) ((65), SI). While our focus was bacterial body shape, flagellum shape also affects swimming performance. Since there is a profound scarcity of detailed data on flagellar morphology, we chose to pair each body shape with its optimal helical flagellum (i.e., optimal *a*, λ, and *n_λ_*: Fig. 2C, SI) to reduce artificial biases in our results, despite this task requiring > 100,000 CPU hours.

#### Chemotactic SNR

Chemotactic SNR is proportional to 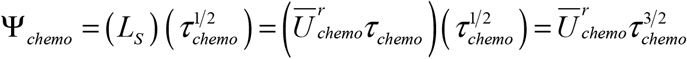 where *L_S_* is the distance between samples, *τ_chemo_* is the sampling time or timescale for loss of orientation, and 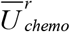 is the effective swimming speed, rescaled such that all equal-volume cells dissipate the same power (SI) (18). While 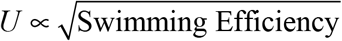 generally (SI), because Swimming Efficiency and Chemotactic SNR are optimized by different flagellum shapes (Fig. 2C, SI), here we again found the optimal helical flagellum shape for each body shape to remove any arbitrary biases in our results (SI). The variation in 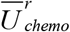 here (Fig. S 6) is thus qualitatively different than the topology of our Swimming Efficiency landscape (Fig. 3A). Since Chemotactic SNR appears to increase without bound with flagellum length (Fig. S 13), flagellar arclength was constrained to a maximum of 15 μm (approximately 24 body lengths of our spherical bacterium) for these optimizations, consistent with observations of *E. coli* (66).

#### Construction Ease

Total absolute Gaussian curvature is defined as 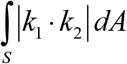 where *S* is the body surface. The absolute value causes local regions of both concave and convex surface curvature to add to total cost. Since cell membrane and wall raw materials also likely incur energetic and material costs, we multiplied this cost function by total surface area *A*, taking the square root to obtain a linear scaling of total cost versus area: 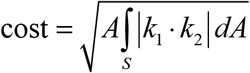. Finally, to facilitate comparison with other tasks and simplify the forthcoming analysis, we inverted the cost function to yield a Construction Ease under selective pressure to be maximized: 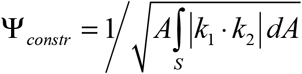. We also considered many other alternative cost functions, detailed in SI, but this one was most consistent with observed bacterial shapes.

### Pareto optimality and Goodness of Fit

For each putative set of evolutionarily crucial tasks (out of 148 combinations), we sampled the interpolant of each task’s performance landscape throughout a discretized grid of 125 x 100 points in morphospace, and removed the dominated points which are beaten at all tasks by at least one other point (full details in SI). For each set of tasks, we computed Hargrove et al.’s (67) goodness of fit (GoF) parameter between the resulting Pareto-optimal region of morphospace and the boundary around observed species shapes as 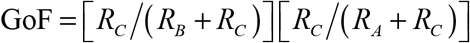 where *R_A_* is the area within the observations but outside the optimal region, *R_B_* is the area within the optimal region but outside the observations, and *R_C_* is the area shared by both the observations and optimal region (Fig. S 15).

### Code availability

The computer code used in this work is available at https://github.com/rschuech/RSBEM.git under the GPL 3.0 license; note that it is under active development.

### Data availability statement

The authors declare that the relevant data supporting the findings of this study are available as supplementary data (Tables S2, S3). Any additional data that support the findings of this study are available from the corresponding author upon request.

## Notes

* We use this term in favor of the well-known “fitness landscape” concept (27) to distinguish performance in individual tasks from overall evolutionary fitness

† For brevity and accessibility, instead of the typical mathematical meaning, we use the term “curvature” loosely to describe the dimensionless fraction of a full circumference formed by the cell centerline

‡ Actually, extremely curved ring-shaped bacterial species *do* exist (e.g. *Thioalkalimicrobium cyclicum* (30), *Rhodocyclus purpureus* (31), *Microcyclus spp.* (28), *Cyclobacterium sediminis* (32)) but none are motile except perhaps “occasional” individuals of *Nitrosovibrio tenuis* (33); we restricted our study to motile bacteria, although the question of why some non-motile bacteria are curved (e.g. *Pelagibacter ubique*) is equally intriguing (1).

§ The keen reader might notice relative maxima in Construction Ease versus *ℒ* for high *𝒦* rods. This is due to increasing *ℒ* causing a penalty from increased surface area, but also a cost reduction via an increase in axial radius of curvature (i.e. a less sharply curving cell centerline); the latter can overtake the former in some regions of morphospace.

** The next-best GoF corresponded to tradeoffs between Tumbling Ease, Chemotactic SNR, and Construction Ease, but we believe this fit (Fig. S 15C) sufficiently worse than the best (Fig. 4) to rule out this alternative hypothesis. Sets of more than 3 tasks (e.g. adding Nutrient Uptake) did not result in improved fits to the data (Fig. S 15, SI).

## Acknowledgments

We thank Oscar Guadayol, Fouad El Baidouri, Mathew Walker, Sei Suzuki, and Chris Venditti for critical feedback.

## Funding

This work was funded by a Leverhulme Trust Research Leadership Award (RL-2012-022) to S.H.

## Author contributions

R.S. and S.H. designed the study, R.S. developed the methodology and software, D.S. provided mathematical guidance, T.H. and R.S. collected morphological survey data and analyzed images. R.S. and S.H. wrote the paper. All the authors discussed the results and commented on the manuscript.

## Competing interests

Authors declare no competing interests.

## Supplementary Information

### Detailed Methods

#### Survey of observed shapes

Micrographs of curved bacteria were collected from the literature as described in Methods of the main text. In a single case (*Vibrio cholerae*), several images were downloaded from an academic website (http://remf.dartmouth.edu/images/), but all other species images were extracted from peer-reviewed publication PDF files (Table S2) via Adobe Acrobat^®^. Species that appeared in images only as spheres or straight rods were omitted. Hence, while we included some straight rod individuals in the dataset, our focus was on curvature since straight rods have already been surveyed elsewhere (1). Images frequently contained a variety of other shapes (e.g. spirals), but only the individuals with shapes representable in our 2D curved rod parameter space (i.e. curved and straight rods, and spheres) were included in our measurements. Images with fewer than 50% of cells exhibiting representable shapes were discarded. Efforts were made to exclude cells that appeared to be undergoing division or sporulation. In transmission and scanning electron microscope images, effort was made to only include cells transected through their centerlines or oriented such that their centerlines were aligned with the image plane.

Cells were segmented using MicrobeJ (2). Due to poor contrast and insufficient resolution, many images required manual pre-processing to exclude cells not fitting the above criteria and to assist the successful segmentation of others. The “Rod-Shaped” shape type was used, allowing for curvature of the cell centerline and outputting its average value as well as centerline arclength and cell width. Scale bars, when present, were used to calibrate measurements and compute equivalent spherical diameter as a measure of size; when scale bars were absent, mean cell width and length values from the article text were used to estimate median size.

To calculate Goodness of Fit (GoF) between the theoretically Pareto-optimal regions and observed bacterial shapes, we computed a polygonal boundary around the latter data points using MATLAB’s *boundary* function with the default shrink factor of 0.5 (Fig. S 1). This is slightly different than the convex hull since it is allowed to envelope the points more tightly. We implicitly included the *ℒ* axis (i.e., all straight rods) as data points in this calculation since previous work (1) has shown the wide diversity of straight rod aspect ratios. While the majority of curved rod species medians exhibited *ℒ* < 10, there were a few highly elongated outliers (Fig. S 1): *Helicobacter mastomyrinus* (*ℒ* = 20.7) and *Methanospirillum hungatei* (*ℒ* = 12.4). Since our simulated morphospace only extends to *ℒ* = 10, we intersected the boundary curve with a vertical line at this location to form a finite-size two-dimensional region (Fig. S 1) that we could compare with our Pareto-optimal regions and compute GoFs.

While size is a critically important trait in bacteria (3), we only considered shape here. This is largely because within our modeling framework, including size as another trait in addition to *ℒ* and *𝒦* would not have any effect on our predictions of Pareto-optimal shapes. Our definition of Swimming Efficiency is scale-invariant (within the bounds of Stokes flow), and similarly for our other tasks (Chemotactic SNR, Construction Ease, Tumbling Ease, Nutrient Uptake), an increase in size would affect all bacterial shapes equivalently. Therefore, a change in size would scale our performances up or down but not change the landscape topology (i.e. shape of contours) – the effects of shape on task performances and Pareto-optimality would be unchanged. This is not to say that there are no interactions between size and shape in bacteria.

For instance, in the case of chemotaxis, one might expect a cost of elongation related to diffusion time for signaling molecules to get between poles of the cell for bacteria with flagella at both poles (4). However, we did not consider this here as it suggests a hard-limit on cell length rather than a continuous tradeoff.

#### Governing equations and the Boundary Element Method

Bacteria live in a fluid environment dominated by the effects of viscosity, with inertia of virtually no importance. This fact is quantified by the Reynolds number (Re = *ρUL /μ*, where *ρ* is fluid density, *μ* dynamic viscosity, *U* swimming speed, and *L* cell size), which is approximately 10^-8^ for a 1 μm bacterium moving at 10 μm s^-1^ in water of density 10^3^ kg m^-3^ and viscosity 10^-3^ Pa s. Since Re << 1, the flow around swimming bacteria is entirely in the creeping flow regime, and the Navier-Stokes equations that govern fluid flow are greatly simplified since the non-linear inertial terms are negligible. The flow of water around a bacterium is thus well-approximated by the Stokes flow equations:

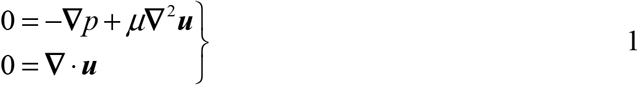

where *p* is pressure, *μ* dynamic viscosity, and ***u*** velocity. These flows are quasi-steady in that the only time dependence is due to time-varying boundary conditions. The Boundary Element Method (BEM) has been extensively used to solve these equations for fluid flows around small particles and microorganisms and can be very efficient due to the effective reduction in dimensionality of the problem from 3D to 2D. In a BEM, velocities anywhere in the fluid domain are related to traction (i.e., surface stress) on the domain boundaries through a boundary integral equation:

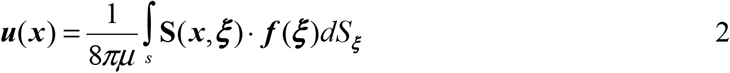

where ***u***(***x***) is the velocity at point ***x***, *S* represents the domain boundaries, ***f*** (***ξ***) is the traction (i.e., the force per unit area imparted by the solid boundary on the fluid) at point ***ξ*** on the boundaries, and, in a traditional BEM, **S**(***x***, ***ξ***) represents the fundamental solution to equation 1, i.e. the flow field due to a singular point force at ***ξ*** termed the “Stokeslet” (see e.g. (5). The presence of the singular Stokeslet in most BEMs introduces a significant disadvantage: the boundary integrals in traditional BEMs not only require high computational effort, but often special treatment by the programmer to be computed at all.

#### The regularized Stokeslet method

The method of regularized Stokeslets (RSM) introduced by Cortez et al. (6) largely ameliorates the burden of programming a BEM. Briefly, Cortez et al. replaced the Stokeslets of the traditional BEM by “regularized Stokeslets,” resembling spatially concentrated but finite-magnitude regions of force density, with the sharpness controlled by the regularization parameter *ε*:

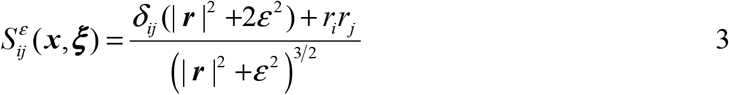

where *r_i_* = *x_i_* − *ξ_i_* and *δ_ij_* denotes the Kronecker delta tensor. This results in the regularized version of equation 2:

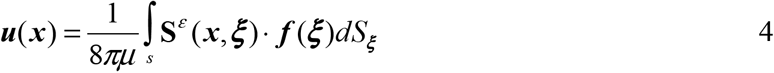

As *ε* is decreased toward zero, 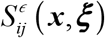 approaches the singular Stokeslet. The boundary integrals arising from the regularized Stokeslet method, although still often near-singular, are readily handled by general purpose numerical integration routines, and the numerical error introduced by a non-zero *ε* can be understood and controlled through convergence studies.

While Cortez et al. introduced an excellent tool for the study of low-Re flows, their original implementation, which employs the same discretization for both traction and numerical quadrature, is too computationally expensive for large-scale studies. Smith (5) later suggested that traction be discretized independently of the quadrature nodes, which is highly advantageous since traction can usually be well represented by far fewer mesh nodes than the number of quadrature points required to accurately compute the near-singular boundary integrals. We refer the reader to Smith (5) and Cortez et al. (6) for full details of this boundary element regularized Stokeslet method, and focus here on differences between Smith’s implementation and ours.

#### Our implementation

While Smith’s proof-of-concept demonstration (5) used a constant-force discretization for traction and a first-order flat triangular mesh, the method presented here uses isoparametric 2^nd^ order (quadratic) triangles as in Shum et al. (7). Although a custom meshing algorithm could have been designed to mesh the curved rod bodies and helical flagella, we desired to maintain maximum generality in the interests of future work and opted to use an existing software suite designed specifically for this task. The Salome Platform (8) includes a geometry creation module as well as a meshing module, and is scriptable via Python to automate the construction and meshing of arbitrary geometries. We used the Netgen (9) 1D-2D algorithm that is included within Salome to construct 6-node (T6) quadratic triangular surface meshes.

Due to the use of adaptive integration for the boundary integrals, the dependence of the solution on the surface mesh refinement was much reduced. In fact, we found that Salome/Netgen could typically only successfully produce fairly refined meshes (Fig. S 2), beyond what would be necessary for accurate simulations. Mesh refinement was controlled primarily by minimum and maximum element size, and to account for bodies of different elongation, these were scaled to be proportional to the square root of body radius. While this strategy worked in most cases, occasional meshing failures were overcome by increasing mesh refinement further. All body meshes passed an automated series of tests (e.g., enclosed volume acceptably close to expected, maximum angle between adjacent element normals sufficiently small) and each body mesh was also visually inspected. Flagella mesh refinement was also controlled by minimum and maximum element size. Meshing of flagella was extremely reliable since the key parameter, flagellum radius, was kept constant for all simulations.

We further improved upon the methods of both Smith (5) and Shum et al. (7) by taking advantage of an existing adaptive quadrature routine for simplices, *ADISMP* (10), to handle the boundary integrals in a highly accurate yet efficient way. We used *ADISMP* to adaptively subdivide mesh elements into smaller triangles until an absolute integration error estimate was below the desired tolerance; this tolerance was scaled with element surface area so that a tolerance per unit element area was effectively used. We placed no limit on the number of element subdivisions allowed, and achieved the fastest convergence with a 5th order integration rule, though higher order rules are included in *ADISMP*.

To take advantage of modern multi-core computer architectures, much of the code was parallelized via the *parfor* construct in MATLAB. In particular, boundary integrals were parallelized over the collocation points (i.e., mesh vertices). In addition, computationally intensive parts of the code (primarily the boundary integrals) were further sped up by approximately a factor of 30 by converting the MATLAB code into compiled C code, with parallelization preserved through OpenMP, via the MATLAB Coder toolbox. Although our MATLAB implementation is limited to shared memory parallelization and cannot take advantage of distributed memory parallelism over multiple compute nodes, reasonable run times (O(1) - O(10) minutes) were observed for our problems on 20-core Intel E6500 servers. Furthermore, sweeps over our *ℒ*, *𝒦* parameter space could be run independently, in parallel, on several servers at once.

#### Model geometry

##### Body

Body geometry was defined as described in Methods and Fig. 1. Note that because we are focused on shape and not size, all of our performance predictions (e.g., Fig. 3) are presented as ratios normalized to the performance of an equal-volume spherical body. Hence the choice of 1 μm^3^ volume for all body shapes is arbitrary though nonetheless realistic for many bacteria.

As part of our parameter space, we included important degenerate cases of a general curved rod. *ℒ* = 1, *𝒦* = 0 is a spherical body. *ℒ* > 1, *𝒦* = 0 yields straight rods (i.e., capsules) of increasing slenderness as *ℒ* is increased. *𝒦* = 1 would theoretically be a ring-like shape with touching poles, but this extreme case is not studied here. In fact, there are two regions of the theoretical morphospace that result in problematic ring-like geometries. As *𝒦* approaches unity particularly for large *ℒ*, it becomes increasingly likely that the flagellum intersects the body as it rotates, and these cases were handled by shifting the flagellum position slightly (details below). In addition, as *𝒦* approaches unity for small *ℒ*, the torus-shaped body eventually becomes self-intersecting as the inner radius approaches zero. Hence, the simulated morphospace is not a full rectangular region (Fig. 2A).

##### Flagellum

The flagellum was modeled as a modified right-handed helix of finite circular cross section (Fig. S 2) as in Shum et al. (7). We note that while most bacteria (e.g. *Escherichia coli, Salmonella typhimurium*) seem to have left-handed flagella that primarily rotate counterclockwise (11, 12), many exceptions of right-handed flagella and clockwise rotation are known (13–17), with the choice being arbitrary for the purposes of this study. Flagellum centerline coordinates are defined according to Higdon’s (18) original suggestion:

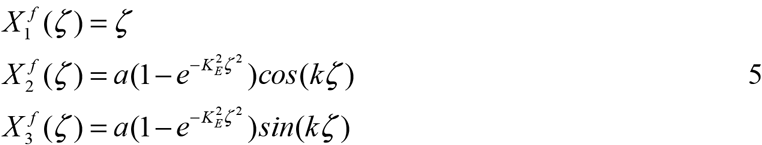

where *a* is the helix amplitude, *k* = 2*π/λ* is the wavenumber and *λ* the wavelength, and *K_E_* is a growth parameter controlling how quickly the helix amplitude increases from zero to *a* as one travels away from the body. The latter modification allows more natural “attachment” of the flagellum to the body, along the helix centerline. The parameter *ζ* varies from zero to *n_λ_λ*, the length of the flagellum axis. As in Shum et al. (7), the flagellum is capped by hemispheres at both ends. The radius of the flagellum was kept constant at 5% of the equivalent spherical radius of the body, resulting in a flagellum radius of 0.03 μm. This is slightly larger than the 1 – 4% typically found in bacteria, but reduces computational cost and is identical to the choice of Shum et al. (7) to facilitate comparison of results. Since the flagella of multiflagellated species often form a bundle during a “run” that is well approximated by a single thick helical tube (19), our choice can also be regarded as an intermediate approximation to both monotrichous and multitrichous bacteria.

Fujita and Kawai (20) found a relatively weak relationship between the amplitude growth rate parameter *K_E_* and swimming efficiency so, like Shum et al. (7), we fixed *K_E_* to vary with flagellum wavelength such that *k /K_E_* = 1. Hence, we varied three independent parameters of our model helical flagella: amplitude *a*, wavelength λ, and number of wavelengths *n_λ_*.

For all simulations, the flagellum was joined near one pole of the body and oriented such that the centerline of the helix was aligned with the centerline of the body (Fig. S 2). As in Shum et al. (7), a small gap equal to one flagellum radius usually existed between the flagellum and body to prevent numerical problems due to the counter-rotation of the two surfaces. However, for ring-like body shapes of *𝒦* near unity, this configuration can lead to intersection of the flagellum with the other pole of the body. Before the simulation started, when any intersections over a flagellum rotation were detected, the flagellum was iteratively shifted back slightly (by half a flagellum radius) until there was enough clearance. This modification allowed such body shapes to be paired with physically realistic flagellum shapes. In all cases the flagellum was allowed to rotate around its axis relative to the body, but otherwise constrained to move with the body.

#### Free-swimming constraints

To simulate the model bacterium’s swimming kinematics, we follow Shum et al. (7) and assume that the body has translational velocity ***U*** and rotational velocity ***Ω***^*B*^ in a body-attached reference frame at any instant in time. Although all points in the body have the same ***U***, a reference point ***x***_*r*_ is needed to define ***Ω***^*B*^; we chose ***x***_*r*_ to be the center of the sphere capping the body-end of the flagellum (i.e., the origin of our body-frame coordinate system, Fig. S 2). The flagellum moves with the body at ***U*** and ***Ω***^*B*^ but additionally rotates relative to the body at rotational speed ***Ω***^*F*^ around the flagellum axis ***e***^*F*^, which is identical to our x-axis (note our definition of the unit vector ***e***^*F*^ is opposite in direction to that of Shum et al.’s ***e***^*T*^). If the relative position of any point ***x*** relative to ***x****_r_* is given as ***x̃*** = ***x*** − ***x***_*r*_, the velocity anywhere on the bacterium’s surface can be written as:

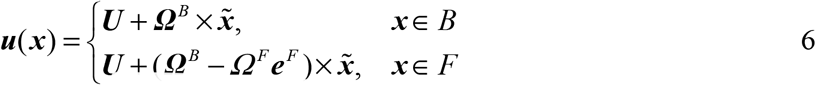

where *B* is the body surface and *F* is the flagellum surface. We have just introduced three new unknowns, the vectors ***U*** and ***Ω**^B^* and scalar ***Ω***^*F*^, in addition to the unknown traction ***f*** on the bacterium in equation 4, so three additional constraints are required to solve the problem.

Since we wish to model a bacterium swimming freely without any externally imposed forces (e.g., gravity is neglected), and since inertia is ignored in the Stokes regime, the sum of both external forces and torques on the bacterium (i.e., the body and flagellum combined) must equal zero at any time. Since the flagellum is constrained to only rotate around ***e***^*F*^ relative to the body, we can also specify the torque imposed by the flagellar motor, which acts between the body and flagellum. These conditions are expressed by the following three constraints:

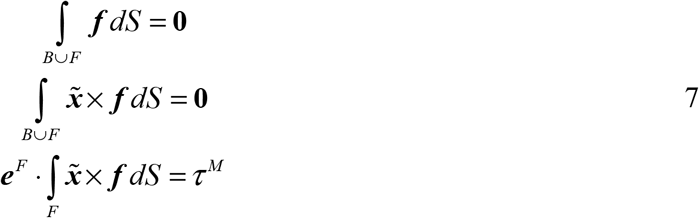

where *τ^M^* is the specified, constant torque applied by the flagellar motor (arbitrarily 10^-6^ N μm for all simulations since this yielded swimming speeds of approximately 20 μm s^-1^). While we used a constant-torque constraint for the flagellar motor, a constant-rotation-rate condition is also commonly applied. We chose a constant torque condition because the torque-rotation rate relationship for most bacterial flagellar motors exhibits an extended plateau region of constant torque independent of rotation rate, with this torque value suggested to be biochemically optimal for a given motor (21–23). We note that many species do not normally operate in the constant-torque region, but in a higher frequency regime where torque linearly decreases with rotation rate (23). Regardless, the assumed torque-speed relationship is irrelevant to our calculation of Swimming Efficiency and our other task performances because this kinematic constraint will merely modify the temporal properties of the swimming trajectory, not its shape nor the energy required to traverse a given distance.

After discretization of the bacterium surfaces into *N* collocation points (i.e., the vertices of the T6 meshes), there are 3*N* unknown traction components and seven unknown kinematic components (the vectors ***U*** and ***Ω***^*B*^ and scalar ***Ω***^*F*^). 3*N* linear equations involving ***f***, ***U***, ***Ω***^*B*^ and ***Ω***^*F*^ can be written by inserting equation 6 into the boundary integral equation (4). Combining these 3*N* equations with the seven equations contained in the free-swimming constraints (7), we can solve for both traction as well as movement kinematics at any instant in time. A dense matrix of equations is obtained, which we solve using MATLAB’s built-in *mldivide* function.

#### Swimming trajectories

Simulation of a freely swimming bacterium requires solving a system of ordinary differential equations (ODEs) to obtain the location ***X*** (*t*) and orientation ***θ***^*B*^ (*t*) of the bacterium in a fixed reference frame as a function of time. In the case of the constant-torque motor condition applied in this study, the phase angle of the flagellum *θ^F^* (*t*) is additionally computed. ODEs (given below) relate the rates of change ***Ẋ***, ***θ̇****B*, and *θ̇^F^* to the instantaneous kinematic variables ***U***, ***Ω***^*B*^, and *Ω^F^*.

##### Adaptive Fourier interpolation

***U***, ***Ω***^*B*^, and *Ω^F^* are required as functions of time, but they depend only on the instantaneous geometric configuration of the body and flagellum. While these kinematic quantities may be updated during each timestep by rebuilding the entire matrix of boundary integral equations and solving them, this is very inefficient for the present problem. This is because the cyclic nature of a single bacterium swimming in an unbounded domain causes ***U***, ***Ω***^*B*^, and *Ω^F^* to be periodic over each revolution of the flagellum. Therefore, interpolants for these functions over flagellar phase angle were created and stored for very fast evaluation at any *θ^F^* during time-stepping.

Trigonometric interpolation via the Fast Fourier Transform (FFT) (24) was used due to rapid convergence for periodic data. Four *θ^F^* evaluation points were used for all free-swimming simulations, since using eight points yielded O(0.001%) difference in effective swimming speed in tests of several different body and flagellum shapes. Typical Fourier interpolants are shown in Fig. S 3.

##### Performance improvement via tensor rotations

For free-swimming problems involving at least one rigid object that may translate and rotate but not deform over time, one may save a possibly significant amount of computational effort by recycling some of the boundary integrals that have already been computed at previous times. Specifically, the individual boundary integrals in equation 2 corresponding to collocation points *and* mesh elements that are both on the same rigid object do not need to be recomputed every time the object translates and/or rotates. This is because translation will have no effect on these integrals, and the effects of rotation can be dealt with relatively quickly via a standard rotation of the tensors that were obtained by integrating **S**(***x***, ***ξ***) previously (i.e. **σ**_*r*_ = **R σ R***^T^* where **σ** is a rank two tensor, **R** is a rotation matrix, and **σ***_r_* is the rotated tensor). Therefore, the entire matrix of equations was only built once for each simulation, when evaluating the initial phase angle *θ^F^* = 0. For other phase angles, only the “interaction” entries of the matrix corresponding to boundary integrals involving collocation points on the body and mesh elements on the flagellum, or collocation points on the flagellum and mesh elements on the body, were recomputed. The integrals for collocation points and elements both on the body were copied unchanged, and the integrals for collocation points and elements both on the flagellum were recycled via tensor rotations with **R** representing a rotation of *θ^F^* around the flagellum axis. (The matrix entries arising from the free-swimming constraints in equation 7 were also recomputed but the computational cost was insignificant). Results of simulations using this shortcut were compared to those without and were confirmed to be identical.

##### Free-swimming ordinary differential equations

To compute the free-swimming trajectory, we followed the general approaches of Keller and Rubinow (25) and Ramia et al. (26), briefly reviewed here. Let ***r***_*B*_(*t*) represent any position vector on the body in the body reference frame, and ***R***_*B*_(*t*) be the same point represented in the fixed reference frame, at time *t*. The transformation from ***r***_*B*_(*t*) to ***R***_*B*_(*t*) consists of a translation by ***X*** (*t*) and a rotation by **A**^−1^ (*t*):

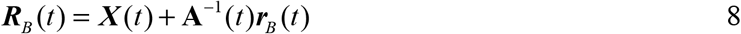

where **A**^−1^ (*t*) is a 3 × 3 rotation matrix (27) expressed in terms of three angles, representing intrinsic rotations around the z, y’, x’’ intermediate axes by 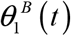,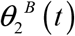,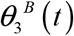, or equivalently, extrinsic rotations around the x, y, z axes by 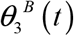,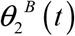,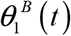 (note, a documented (28) typographical error exists in (27), corrected here):

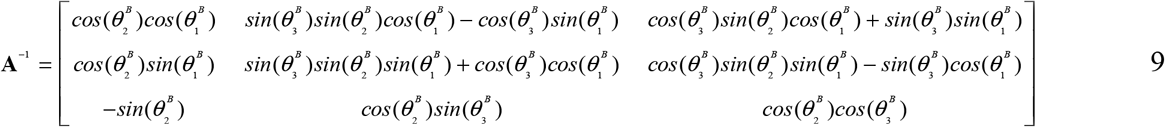

The rate of change of ***X*** (*t*), ***Ẋ***, is related to **A**^−1^ (*t*) and the body frame velocity ***U*** (*t*):

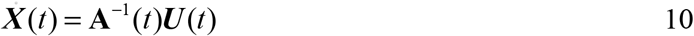

The angles ***θ***^*B*^ and their rates of change ***θ̇***^*B*^ in the fixed frame are related to the bacterium’s angular velocity ***Ω***^*B*^ in the body frame (27):

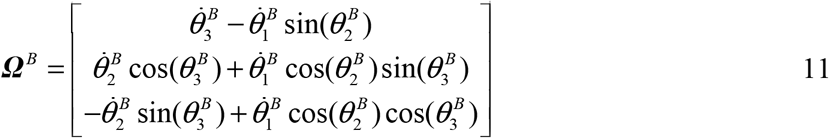

These equations can be solved for ***θ̇***^*B*^ in terms of ***Ω***^*B*^ and ***θ***^*B*^:

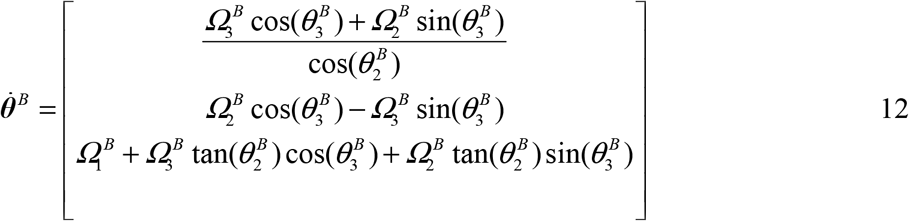

Note that to specify rotations, we use Goldstein et al.’s (27) “xyz-convention” (Tait-Bryan angles) instead of the “x-convention” (Euler angles) used by Ramia et al. (26), so that the singularity in ***θ̇***^*B*^ occurs at 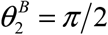 instead of 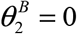 and all components of ***θ***^*B*^ can be initialized to zero. Conveniently, 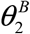 did not approach π/2 in any of our simulations and special treatment of the singularity was not needed.

The position of any point on the flagellum over time obeys expressions identical to equations 8 and 9 except that 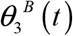 is replaced by 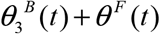 to account for the rotation of the flagellum relative to the body. The rate of change of the phase angle of the flagellum is trivially identical to *Ω^F^* since both are defined in the body frame:

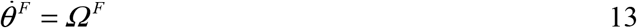

##### Numerical solution

MATLAB’s *ode45* was used to solve the system of differential equations 10, 12, and 13 for the evolution of bacterial position ***X***, orientation ***θ***^*B*^, and flagellar phase angle *θ^F^* over time. For all simulations, the midpoint of the body centerline ***X***_0_ was used as the initial condition (Fig. S 2) and chosen as a reference point to generate swimming trajectories (Fig. 2B, Movie S1). The initial body orientation angles 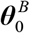 and flagellum phase angle 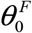 were both set to zero. Time-stepping was continued until an effective average swimming speed could be accurately determined. Since evaluating the Fourier interpolants is computationally cheap, this was done by brute force: a simple linear trajectory parameterized by effective swimming speed *U̅_swim_* was fit to the actual, roughly helical swimming trajectory. The simulated time period was repeatedly doubled until the calculated *U̅_swim_* changed less than 0.05 % between two consecutive doublings. This convergence criterion was achieved within seconds of simulated time for the majority of simulations.

#### Convergence and validation of numerics

Convergence studies were performed on several body-flagellum combinations to determine satisfactory values for the regularization parameter *ε*, the absolute integration error tolerances in *ADISMP*, the *ode45* absolute and relative error tolerances, and the tolerance used to terminate time-stepping when computing *U̅_swim_*. After some preliminary trial and error to determine reasonable starting values, all tolerances were simultaneously halved repeatedly until the maximum change of any key variable (i.e., *τ_chemo_*, *U̅_swim_*, Swimming Efficiency Ψ_*swim*_, and Chemotactic SNR Ψ_*chemo*_ – see next sections) was smaller than 0.01%. Finally, each tolerance parameter was repeatedly doubled one at a time to ensure that computational time was not being wasted on unnecessary refinement of any parameter. In addition, convergence with respect to the surface mesh element size was tested by halving min and max element size tolerances. The meshes were generally over-refined (due to limitations in SALOME (8) and/or Netgen (9)) and the estimated error in the above variables due to the approximation of the geometry by surface meshes was < 0.02%.

We validated the code by testing Shum et al.’s (7) optimal spheroidal body and flagellum combination, and obtained the same swimming efficiency (i.e. their “power efficiency”) to within 1.3%, comparable to their reported level of accuracy. To test our Brownian diffusivity calculations that yield *τ_chemo_* and *τ_tumble_*, we placed a triaxial ellipsoid at an arbitrary location and orientation relative to our coordinate system and calculated the translational and rotational diffusivities. Comparing to the exact solutions for the translational and rotational diffusion coefficients of triaxial ellipsoids (1), the maximum error was < 0.02%. We also recovered the correct “center of diffusion” (i.e., the centroid of the ellipsoid, different than our origin) and the three principal axes of diffusion (i.e., the three axes of the triaxial ellipsoid, different than our coordinate axes).

#### Calculation of evolutionarily relevant tasks

##### Swimming Efficiency and Speed

Swimming Efficiency Ψ_*swim*_ was based on the power dissipated by the rotating flagellum as in (7, 29) and calculated as the ratio of the power required to translate a sphere of the same volume as the body at the effective swimming speed, to the power actually dissipated by the model bacterium in rotating the flagellum:

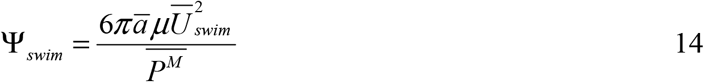

where *ā* is the equivalent spherical radius of the body (approximately 0.62 μm for all bodies in this study), *μ* is dynamic viscosity, *U̅ _swim_* is the effective swimming speed, and *P̅^M^* is the average power dissipated by the flagellar motor, equal to the product of the imposed constant motor torque and the average rotation rate: 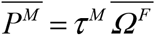. Although this is not a true thermodynamic efficiency, many studies of bacterial swimming have used essentially this same measure of energetic effectiveness. To facilitate interpretation of results, we normalize performances to that of a spherical body when plotting the performance landscape (Fig. 3A): 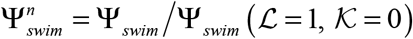

Note that because we assume that the power available to the flagellar motor scales with cell volume, all our constant-volume cells have the same available power (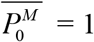 femtoWatt), but this is difficult to enforce *a priori* in simulations. If the simulated swimming kinematics of all cells are rescaled (see *Chemotactic Signal/Noise Ratio*) to enforce this condition, we obtain rescaled swimming speeds 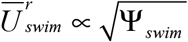 that can be fairly compared across body shapes (e.g., Movie S1). Since this is a simple monotonicity-preserving transformation of Ψ_*swim*_, the performance landscape for *U̅_swim_* would look qualitatively identical to that of 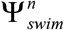 (Fig. 3A) with only the contour values (not topology) differing. Hence there is no need to consider swimming speed separately from swimming efficiency in the subsequent analysis of Pareto optimality.

Shum et al. (7) did consider an alternate efficiency metric based on torque as the limiting quantity instead of power. While this was motivated by the constant-torque region of the torque-speed curve of typical flagellar motors (22, 23, 30), they found that the torque-optimized model bacterium (particularly the flagellum) was of a somewhat unusual shape (e.g. very short), and would require a very high flagellar rotation rate to achieve typically observed swimming speeds. Therefore, we did not consider optimizing bacterial morphology on the basis of torque efficiency.

##### Brownian diffusivities and timescale for loss of orientation (τ_chemo_ and τ_tumble_)

The Brownian motion of highly symmetric particles such as ellipsoids can be fully described by three translational diffusivities and three rotational diffusivities, one for each axis, where the axes correspond to the axes of symmetry (31). However, general particles with little or no symmetry (e.g. curved rods with helical flagella) tend to have at least some coupling between translational and rotational motions, and so require a much more involved treatment to obtain the correct Brownian diffusion parameters. We followed the algorithm recommended by Wegener (32): translational, rotational, and coupling friction coefficient matrices (Wegener’s ^*t*^**K**, ^*r*^**K**_*P*_, and ^*c*^**K**_*P*_, respectively) were determined by simulating forced translation and rotation of the geometry (i.e., body + flagellum for Chemotactic SNR Ψ_*chemo*_, body only for Tumbling Ease Ψ_*tumble*_) along / around the x, y, and z axes. In the body + flagellum case, we neglected the effect of flagellum phase angle after testing showed this to be very small for the geometries of interest here. These forced simulations were done similarly to the force-free flow simulations described above, except surface velocities ***u***(***x***) in equation 4 were directly specified, and the kinematic constraint equations (7) were omitted along with the free-swimming unknowns ***U***, Ω^*F*^, and Ω^*F*^. (Note that for these six forced-flow simulations, matrix assembly only needs to be performed once, and then the system solved once for six different right-hand sides. Furthermore, the coefficients matrix can be almost completely recycled for a subsequent free-swimming simulation by simply concatenating on the kinematic variables and constraints.) Wegener’s (32) process was then followed closely with one exception: while ^*t*^**K** and ^*r*^**K**_*P*_, and consequently the translational and rotational diffusion matrices ^*t*^**D** and ^*r*^**D**_*D*_, should be symmetric matrices for any arbitrarily shaped particle (32), they are not exactly symmetric here due to numerical errors. If used as-is, this would result in the three principal axes of diffusion not being exactly orthogonal (the angles tending to be a few degrees off from 90°). We chose to eliminate this small error by “symmetrizing” ^*t*^**D** and ^*r*^**D**_*D*_ according to the following algorithm (shown as MATLAB code) for generic square matrix **M**:

`[Eigenvectors_0, Eigenvalues_0] = eig(M)`
`M_sym = Eigenvectors_0 ∗ M ∗ (Eigenvectors_0)’`
`[Eigenvectors, Eigenvalues] = eig(M_sym)`

After symmetrizing, the final eigenvectors are orthogonal to each other to within machine precision. These eigenvectors represent the principal axes for translational (in the case of **M** = ^*t*^**D**) and rotational (in the case of **M** = ^*r*^**D**_*D*_) diffusion (the latter shown in Fig. 2B).

Finally, following Dusenbery (1), we required the timescale of which loss of orientation of the swimming direction occurs due to Brownian rotation (*τ_chemo_* and *τ_tumble_*; we frequently drop the distinction and subscripts in the following for brevity). Since rotations around the swimming axis will not change the direction of this axis, *τ* only depends on the rotational diffusion coefficients for the two perpendicular axes (Fig. 2B). In Dusenbery’s analysis of ellipsoidal bodies, the assumed swimming axis was the same as one of the principal axes of diffusion. However, this is not necessarily the case for asymmetric body shapes like curved rods. In an effort to mitigate this discrepancy, we first rotated the rotational diffusion matrix ^*r*^**D**_*D*_ to align one of the principal axes (the axis closest to the flagellum centerline for *τ_chemo_*, or the axis closest to the vector from rear to front body pole for *τ_tumble_*) with the average swimming direction, yielding 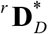. However, due to the rotated coordinate system, off-diagonal coupling terms now appear in 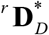, causing the diagonal entries and thus *τ* to depend on the arbitrary rotation of the 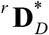 principal axes *around* the swimming axis. Therefore, we computed 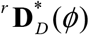 for all possible values of this arbitrary rotation angle *ϕ*, and following Dusenbery (31), calculated 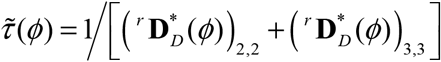 and used the maximum value *τ* = max (*τ̂*(*ϕ*) to define*τ_chemo_* (Fig. S 4) and *τ_tumble_* (Fig. S 5). Although this approach to calculating *τ* is approximate, it avoids resorting to Monte Carlo simulations of swimming combined with stochastic Brownian motion in three dimensions.

While Dusenbery’s approach assumes straight-line swimming paths, in reality the swimming path is typically a helix or more complex periodic trajectory with amplitude and wavelength dependent on the bacterium geometry (Fig. 2B, Movie S1). However, the temporal periods of our simulated trajectories (which matched the body rotation rates around the flagellum axis, Ω_1_^*B*^) were at least an order of magnitude shorter than the predicted *τ_chemo_* and *τ_chemo_.* Therefore, we take this as a post-hoc justification for neglecting the non-linear path shape for the purposes of modeling Brownian rotations.

##### Chemotactic Signal/Noise Ratio

As discussed in the main text, following Dusenbery (31) we considered a temporal chemotaxis strategy in which consecutive samples over time are taken. While spatial sampling strategies using e.g. fore-aft or laterally separated chemosensors on the cell are theoretically possible (33, 34), the observed clustering of chemoreceptor proteins at a single pole of species with a single polar flagellum (35, 36) is evidence against a spatial sampling mechanism, and we did not consider it further.

We defined a quantity proportional to Chemotactic SNR via a temporal comparison mechanism as 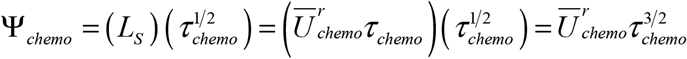 where *L_S_* is the distance between samples, *τ_chemo_* is the timescale for loss of orientation of the swimming direction due to rotational diffusion, and 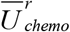 is a rescaled effective swimming speed (discussed further shortly); the other parameters affecting SNR are environmental (31) and thus considered to be constant across our simulations.

Since Ψ_*swim*_ and Ψ_*chemo*_ are optimized by different flagellum shapes (see *Flagellar shape optimization* below), the flagella paired with each body shape are different between these two tasks, and so are the swimming performances (e.g. *U̅ _swim_* vs *U̅ _chemo_*). Thus, the swimming speeds used to calculate Ψ_*chemo*_ are not simply proportional to 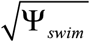, but are determined from new simulations identical to those performed for Ψ_*swim*_ except with different flagella shapes. To compare across different body shapes, we rescaled the simulated swimming speeds *U̅_chemo_* used to calculate Ψ_*chemo*_:

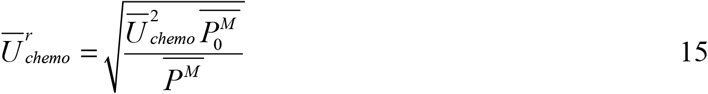

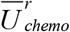 represents the rescaled effective swimming speed, if the average power dissipated by the flagellar motor for Ψ_*chemo*_ calculations were 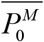 instead of the morphology-dependent 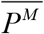 originally realized in the simulations. This rescaling essentially assumes constant power density for all our equal-volume cells. Here, we chose 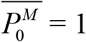 femtoWatt, since this yields realistic swimming speeds of about 20 μm s^-1^ over much of our *ℒ*, *𝒦* parameter space. However, this choice is arbitrary and cancels out when we normalize Ψ_*chemo*_ to that of a spherical body, yielding 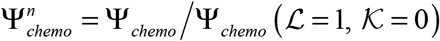.

While the performance landscape of 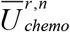 (Fig. S 6) is qualitatively different than the 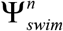 landscape (Fig. 3A), a tradeoff between 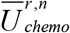 (which, like 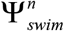, favors curved rods) and 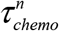 (which favors straight rods, Fig. S 4) is still apparent and leads to the performance landscape of 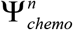 (Fig. 3B) discussed in the main text.

##### Tumbling

While Chemotactic SNR captures the ability of a bacterium to reliably detect a concentration gradient, it does not fully describe the overall ability of a cell to migrate toward or away from the source. Real bacteria often exhibit movement trajectories characterized by nearly straight runs interspersed with near-stationary reorientation events (37). The latter are useful because they represent the only way a bacterium can purposely alter course (beyond simply reversing or waiting for rotational Brownian motion) if it has determined that it has been swimming in the ‘wrong’ direction. Reorientation behaviors vary among species, with peritrichous bacteria like *E. coli* tending to “tumble” due to the unbundling of the flagella (38), and monotrichous bacteria such as *Vibrio* spp. and others displaying a “flick” due to a buckling instability in the flagellar hook (39, 40). While some progress has been made in modeling the physical dynamics of tumbling (41, 42), these reorientation behaviors are generally very difficult to explicitly simulate. However, the importance of cell shape on the ability to reorient has been documented experimentally: filamentous or artificially elongated *E. coli* are unable to tumble, probably due to increased resistance to rotation around the short axes (43, 44). Therefore, we take a simple approach to modeling the effect of body shape on ease of reorientation: if we assume that all reorientation mechanisms are due to randomly located and oriented perturbing forces applied by the flagella to the body, the overall process is reminiscent of Brownian motion. That is, a body shape with a high Tumbling Ease will tend to quickly lose the orientation of its swimming axis when subjected to random perturbing forces, assuming cells of equal volume expend the same power on active reorientations. Thus, Tumbling Ease is simply defined as Ψ_*tumble*_ = 1/*τ_tumble_* and is therefore diametrically opposed to Ψ_*chemo*_, with elongation having opposite effects, and a sphere intuitively maximizing Ψ_*tumble*_ but minimizing Ψ_*chemo*_. However, here we only include the body, not flagellum, in our calculation of Ψ_*tumble*_ since we are assuming that the flagella themselves are the source of the random perturbating forces. Although this is a simplification of reorientation behaviors, Tumbling Ease roughly quantifies the ease with which different body shapes can undergo these processes.

##### Nutrient Uptake

A fundamental task for all bacteria is the uptake of small molecules such as nutrients, for which diffusion from the bulk fluid to the bacterium surface may be a limiting step (45). It is intuitive that a sphere should have the worst possible performance here, since it has the lowest surface area for a given volume; any departure from spherical should increase diffusive uptake per unit volume (46, 47). Furthermore, in the case of curvature, it is intuitive that ring-shaped rods should be outperformed by straight rods of the same elongation since the latter have the same surface area but do not suffer from any “shielding” of inner surfaces. To quantify this variation with shape, we simulated steady-state diffusive transport of a small molecule (ammonium, molecular diffusivity = 1.64∙10^-5^ cm^2^ s^-1^ (48)) to curved rods immersed in an unbounded domain.

While the boundary element method can also be applied to the diffusion equation (49), we used COMSOL Multiphysics v. 5.1 to perform these simple simulations. Body geometries were exported from SALOME in IGES format and imported into COMSOL via the CAD Import Module; the flagellum was neglected here. The bacterium was centered inside a very large spherical domain (i.e., approximately 60 cell body lengths) with boundary conditions of zero concentration on the bacterium surface and an arbitrary concentration of 1 on the sphere surface (this cancels out when computing performance relative to a spherical body). Unstructured tetrahedral meshes were generated via the software’s native routines. The output variable of interest, Nutrient Uptake Ψ_*uptake*_, was calculated as the surface integral of steady-state diffusive flux of ammonium into the cell. Convergence studies of spherical domain size and tetrahedral mesh refinement were performed to achieve 0.5% error or better in Ψ_*uptake*_. We neglected the effect of advection due to bacterial swimming because the increase in mass transport is predicted to be small for typical bacteria (50, 51); to rigorously test this, we switched to a cubical domain and imposed a free-stream velocity boundary condition of 50 μm s^-1^ (a typical observed bacterial swimming speed) on one face of the domain for several different cell shapes in our morphospace. In all cases, the increase in uptake over the purely diffusive case was at most about 1%, which was substantially smaller than the variation in diffusive uptake across the morphospace (Fig. S 10).

As expected, we find that the optimal shape for 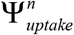 is straight (*𝒦* = 0) and as elongated as possible, while the worst shape is a sphere since it is the most compact. An *ℒ* = 10 straight rod achieves over 50% higher Nutrient Uptake than a sphere. While the effect of diffusive “shielding” due to increasing *𝒦* is much weaker, it nonetheless exerts selective pressure toward lower *𝒦*.

##### Construction Ease

Following Dusenbery (31), we assumed that cells with more sharply curving surfaces are more costly to construct and maintain. In his study of ellipsoidal cell bodies, Dusenbery limited his morphospace to ellipsoids with minimum radius of curvature above some arbitrary, small fraction of the radius of a spherical cell of equal volume. We took this line of reasoning further by defining a cost of construction for any cell shape, based on surface curvature.

At any point on a three-dimensional surface, one can calculate two signed principal curvatures (*k*_1_, *k*_2_), which correspond to the minimum and maximum possible radii of a circle constrained to be tangent to the surface at that point. The signs of *k*_1_ and *k*_2_ depend on whether the surface is locally concave, convex, or saddle-like at that point.

Our generalized curved rod cell shapes are made up of three primitive shapes: the sphere, cylinder, and torus. The principal curvatures for these primitive shapes are:

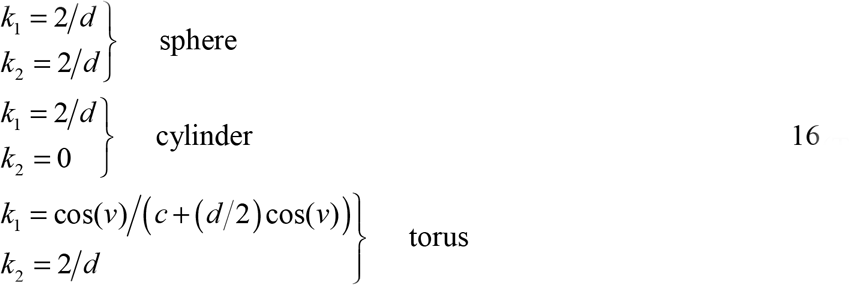

where *d* is the diameter of the sphere or cylinder, or minor diameter of the torus, *c* is the major torus diameter, and *v* is a parameter (0 ≤ *v* ≤ 2*π*) indicating angular position around the torus tube (i.e. inner versus outer).

To arrive at a scalar overall construction cost for a given body shape, one first needs to combine *k*_1_ and *k*_2_ into a single metric at each surface point. We considered three commonly used choices: the Gaussian curvature *K* (not to be confused with dimensionless cell centerline curvature *𝒦*), mean curvature *H*, and maximal absolute curvature *M* (52).

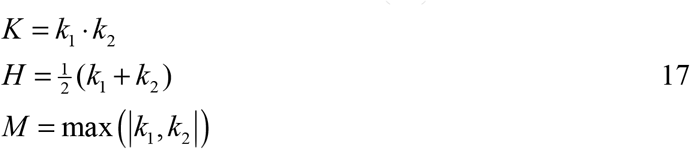

In the case of *K* and *H*, while these standard definitions distinguish between positive and negative curvature, this distinction might not be appropriate when considering costs of cell shape if both types of surface curvature (e.g., invaginations and protrusions) are biologically costly. Therefore, we considered not only these standard metrics of Gaussian and mean curvature, but also absolute variants, noting that in the case of *H* there are two plausible ways of incorporating an absolute value:

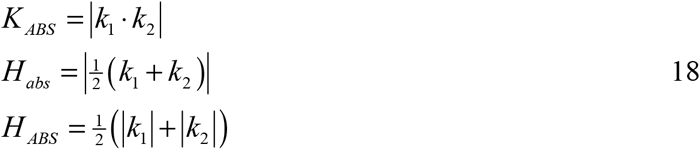

To arrive at a single value describing the overall construction cost of a particular shape, we considered the integral of each of these metrics (*K*, *H*, *M*, *K_ABS_*, *H_abs_*, *H_ABS_*) over the cell surface *S*. We also considered an additional cost component due to cell surface area itself, i.e. the raw material comprising the cell membrane and wall. We thus considered variants of these cost functions that are multiplied by body surface area *A*. Since cost should intuitively scale linearly with surface area (e.g., as in a thought experiment of several identical cells), we took the square root of the area-weighted cost variants to achieve this scaling; this was purely for ideological reasons and does not affect our conclusions since this is a monotonic transformation (see *Pareto optimality*). Lastly, we also considered a very simple cost function equal to surface area alone, in case surface curvature isn’t actually important. To match our convention with other tasks in which higher performance values are better, we inverted all of these putative cost functions to obtain, in total, 13 possible formulations *E* of Construction Ease Ψ_*constr*_:

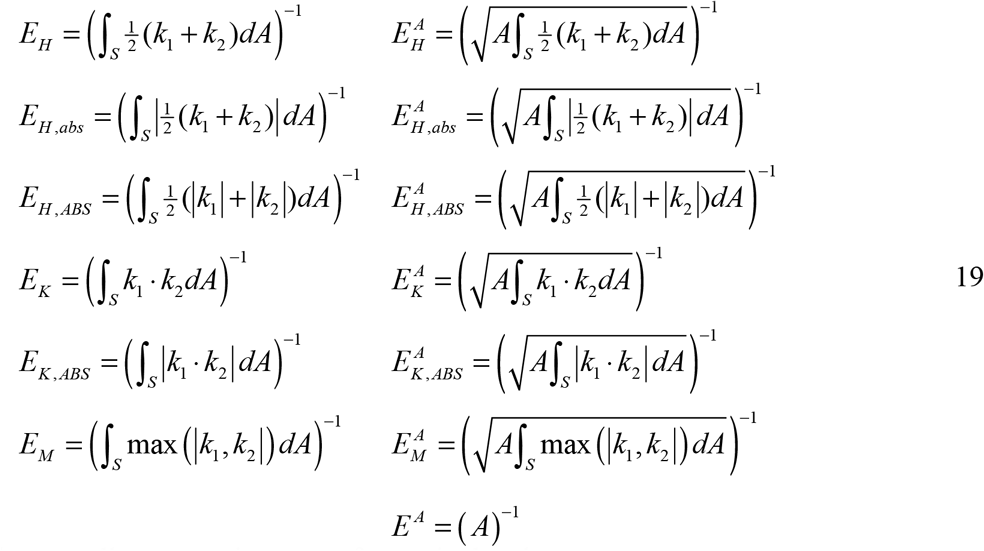

In the case of curved rods with *𝒦* > 0, the integrals in relevant *E* formulations over the toroidal portion of the cell surface were performed numerically using MATLAB’s *integral* function.

In our analysis of Pareto optimality, we considered all 13 *E* formulations by setting Ψ_*constr*_ = *E* for each particular formulation, one at a time. Most of these *E* formulations (equation 19) are dimensioned quantities and thus dependent on size. Therefore, similarly to other tasks, we normalized Construction Ease by a spherical body’s performance to obtain a dimensionless metric dependent only on shape: 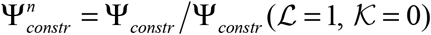.

It is immediately apparent from the Performance Landscapes of the various Construction Ease formulations (Fig. S 11) that some may be of dubious biological realism: *E_H_*, *E_H_* _,*abs*_, *E_K_*, 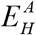, 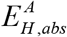 and 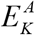 are not maximized by a sphere in contrast to intuition, *E_K_* and *E_K_*_,*ABS*_ are constant for all straight rods regardless of *ℒ*, and *E_M_*, 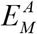, and *E_A_*, are independent of *𝒦* for large regions of morphospace, despite the intuitive hypothesis that increasing *𝒦* should always be costly. This leaves only *E_H_*_, *ABS*_, 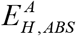, and 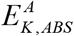 as options that fulfill all intuitive expectations, with 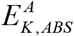 being particularly attractive since the total absolute Gaussian curvature is a dimensionless descriptor of how sharply curved the cell surface is, while *A* accounts for cell size. However, lacking concrete justification for these expectations, we nonetheless generated Pareto-optimal sets of shapes based on all 13 Construction Ease formulations to objectively determine which were most consistent with the observed shape data.

We note that hydrodynamic forces during swimming might tend to deform cells and lead to additional construction costs for highly elongated curved rods, but we did not consider this possibility here.

#### Performance landscapes

To generate performance landscapes, we simulated a grid of body shapes at spacing increments of 0.5 for *ℒ* and 0.05 for *𝒦*. However, because Swimming Efficiency displayed complex contour topology at low *𝒦* and we wanted to accurately locate both the global and local maxima, for this task we refined the grid near these regions (Fig. S 12). And since all Construction Ease formulations were based solely on geometric considerations and did not require any fluid dynamics simulations, they could easily be evaluated at any point in the morphospace, so we computed them over a highly refined grid of 300 x 300 points.

For most tasks, continuous pseudocolor plots (i.e., performance landscapes) were generated by constructing two-dimensional interpolants for the (sometimes scattered) data representing performance for each simulated body shape. We used the *scatteredInterpolant* MATLAB function here with “natural” interpolation. To avoid interpolation artifacts associated with the scaling of morphospace coordinates, we normalized both the *ℒ* and *𝒦* coordinates of each data point to the range [0, 1] when generating the interpolants and then back-transformed after evaluating the interpolants.

#### Flagellar shape optimization

Since we investigated a five-dimensional (5D) parameter space (i.e., *ℒ*, *𝒦* for the body and amplitude *a*, wavelength *λ*, and # wavelengths *n_λ_* for the flagellum), an exhaustive, grid-based sweep over all parameters simultaneously was not feasible. Furthermore, even if we could compute a Pareto-optimal region of 5D morphospace, the paucity of flagella morphology data in the literature would have precluded comparison with observed morphologies in 5D, especially when trying to distinguish the goodness of fit (GoF, see *Goodness of fit*) of Pareto regions resulting from different sets of tasks.

Perhaps the easiest way of dealing with this issue would be to pair each of our body shapes with the same, arbitrarily chosen flagellum shape. However, since the fluid dynamics of swimming are determined by both body and flagellum shape, an arbitrary chosen flagellum would be better suited to some body shapes than others and would therefore introduce a bias in any relevant performance landscapes. Therefore, we eliminated such bias by independently finding the optimal flagellum shape for each simulated body shape, as Shum et al. (7) did for swimming efficiency of spheroidal bodies. Furthermore, since the optimal flagellum shape for Ψ_*swim*_ is different than for Ψ_*chemo*_, we performed these flagellum optimizations separately for each task. While this means that, in a sense, each body in our simulated morphospace simultaneously has two different flagella (one optimized for Ψ_*swim*_, another optimized for Ψ_*chemo*_), our approach eliminates any artificial bias caused by arbitrarily chosen flagellum shapes.

To simultaneously determine the three optimal flagellum parameters (*a*, *λ*, and *n_λ_*) for a given body shape, MATLAB’s *fminsearch* was used. This is a direct search algorithm, which does not rely on derivatives of the objective function. We initially tried faster algorithms such as *fmincon* that utilize approximate derivatives, but due to small numerical errors, the objective functions were too noisy to yield reliable solutions. We first found the optimal flagellum for a body shape near the center of our morphospace; thereafter, we moved radially outward, using the closest known solution as the initial guess for *fminsearch*. The flagellum parameters were centered and scaled to improve algorithm performance. Termination of the optimization algorithm was controlled using TolX = 0.0005. To eliminate small levels of numerical error due to occasional failure of *fminsearch* to find the best solution, we fit 2D thin-plate smoothing splines (MATLAB’s *tpaps* using the default smoothing parameter) to the resulting landscapes of optimized *a*, *λ*, and *n_λ_* and calculated task performance landscapes and Pareto-optimal regions based on these smoothed flagellum shape landscapes.

Ψ_*swim*_ of each body shape in our morphospace is optimized by a finite-sized flagellum. Interestingly, these optimal flagella are all roughly the same shape (i.e. *λ* seems proportional to *a*, and *n_λ_* varies little), and appear to scale with some measure of body size, with the largest flagella paired with the most elongated semicircular rods (Fig. S 14).

While attempting to find optimal flagella shapes for Ψ_*chemo*_, we quickly discovered that this variable appears to increase without bound with flagellum length (i.e., with either *λ* or *n_λ_*, Fig. S 13). Although Ψ_*swim*_ and thus effective speed 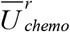 do exhibit maxima relative to flagellum length, the timescale for loss of orientation τ_*chemo*_ appears to increase monotonically and at a faster rate than 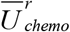 decreases (Fig. S 13). The latter effect dominates, causing Ψ_*chemo*_ to also increase without bound with flagellum length. Given the lack of a finite-sized optimal flagellum for Ψ_*chemo*_, we chose to place a constraint on flagellum arclength ≤ 15 μm for this task, consistent with observations of *E. coli* (53). Since the best flagellum for Ψ_*chemo*_ under this constraint will always be 15 μm long for any body shape, this reduces the dimensionality of the optimization problem for a given body from three to two. Hence, *n_λ_* becomes a (numerically evaluated) function of *a* and *λ*. Although flagellum arclength is fixed, there is still substantial variation in the Ψ_*chemo*_-optimized flagellum shape parameters over our body morphospace (Fig. S 14) due to tradeoffs between τ_*chemo*_ and 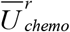 (Fig. S 4, Fig. S 6).

#### Pareto optimality

As discussed by Shoval et al. (54), the space of all possible morphologies (i.e., the morphospace) can be reduced to a set of Pareto-optimal morphologies (called *non-dominated solutions* in other literature) that outperform suboptimal morphologies (i.e., *dominated solutions*) at all tasks that contribute to overall evolutionary fitness. Within the Pareto-optimal set of morphologies, performance in one task can only be improved at the expense of one or more other tasks; the (often extreme) morphologies that each exhibit maximal performance in one task are called the *archetyp*es by Shoval et al.

In their analysis, Shoval et al. assume that the performance landscapes of all relevant tasks are mathematically very well-behaved, with circular contour topology (54). Given these assumptions, they show that the Pareto-optimal region of morphospace resulting from tradeoffs between such tasks is a flat-sided polytope (flat-edged polygon in 2D), with the archetypes for each task occurring at the vertices. If slightly more complicated elliptical contours are assumed, Shoval et al. show that the boundaries of the Pareto-optimal region become “mildly” curving (54). Consequently, by fitting flat-sided polytopes to observed morphotypes of various groups of organisms, Shoval et al. determined what the archetypical morphotypes are. Finally, leveraging abundant data on the ecology of these groups, they inferred the tasks that the archetypes seem to be specialists in, and thus the tasks likely constraining the evolution of morphology within those groups.

We took a different approach than Shoval et al. for two reasons: firstly, the lack of detailed information on the ecology of most bacterial species precludes the inference of tasks based on such information. Secondly and more importantly, in contrast to Shoval et al.’s simplifying assumptions (but see (55)), several of our performance landscapes were rather complicated functions with complex topology (i.e., non-elliptical contours (Fig. 3A,C), relative maxima (Fig. 3A), non-finite archetypes (Fig. 3B)), causing the resulting Pareto-optimal regions unlikely to be simple triangles with archetypes at the vertices. For example, in Fig. 4 we obtain a large two-dimensional optimal region despite the archetypical morphologies that maximize Construction Ease (a sphere, Fig. 3C) and Swimming Efficiency (a slightly elongated pill, Fig. 3A) being very close together in morphospace.

Therefore, we considered many possible sets of tasks, directly computing the resulting Pareto-optimal regions without any assumptions of what they might look like, and finally inferring the most important set of tasks by finding the best fit with observed data. Instead of computing Pareto front edges via points of tangency between contour lines as Shoval et al. discuss, we used a less elegant but more direct and robust method: we sampled the interpolant of each task’s performance landscape throughout a discretization of the morphospace (i.e. a grid of 125 x 100 points, excluding non-physical shapes), and then simply removed the dominated points which are beaten at all tasks by at least one other point, leaving the non-dominated points that comprise a discretized Pareto-optimal region (56). While there are algorithms that aim to directly find Pareto-optimal solution sets without exhaustively evaluating the objective functions over the entire parameter space as we did, in this study we were not only interested in the Pareto-optimal region of morphospace but also the performance in each task over the entire morphospace.

We considered many possible combinations of Swimming Efficiency 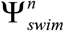, Chemotactic SNR 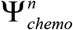, Nutrient Uptake 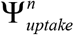, Tumbling Ease 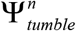, and at most one Construction Ease 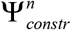 formulation (of 13 putative *E* variants). We considered sets of three, four or five of these putative tasks; we knew *a priori* from Shoval et al. (54) that at least three were needed since, even in the case of complex performance landscape topology, a tradeoff between just two tasks can only result in a 1D Pareto front (i.e., a curve), not a filled 2D region as formed by observed curved bacterial species (Fig. 1). In all, we generated 148 hypothetical Pareto-optimal regions.

An advantage (or disadvantage, depending on context) of Pareto optimality framework is that it is insensitive to the precise magnitudes of each task over the morphospace – in the discrete case employed here, only the rank of each point in the morphospace matters. That is, whether one morphology is 0.1% or 1000% better than another does not matter, only that it is better. Any monotonic transformation can therefore be applied to any task’s performance landscape without affecting which points are Pareto-optimal. This is perhaps most relevant to our consideration of construction cost. For example, we chose to simply invert construction cost to obtain Construction Ease Ψ_*constr*_, but any alternative operation resulting in the same contour topology would be acceptable. Nor does taking the square root of some cost functions to obtain linear scaling with area have any effect on our Pareto optimality results. Likewise, we could have replaced Swimming Efficiency Ψ_*swim*_ with 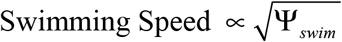 in our sets of putative tasks with no change to the resulting Pareto-optimal regions.

Since the boundaries between optimal and suboptimal regions should be smooth in the absence of any numerical errors, we smoothed the discretized Pareto-optimal regions in Fig. 4 slightly. First we extracted the points along the upper boundary of the main optimal region as well as around the small optimal island using MATLAB’s *bwboundaries*, then fit parameterized splines to the *ℒ* and *𝒦* coordinates of these points. The splines for the main region were constrained (57) to pass through *ℒ* = 1, *𝒦* = 0 (i.e., a sphere, which maximizes 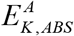 and so must be Pareto-optimal) and have five optimally placed knots, and the splines for the island region were constrained to have four optimally placed knots and periodic end conditions since the boundary is a closed curve in that case. The original, unsmoothed, discretized Pareto-optimal points corresponding to Fig. 4 are shown in Fig. S 15A.

#### Goodness of fit

To quantify how well our theoretical Pareto-optimal shapes matched observed species morphologies, we required a goodness of fit (GoF) parameter. We based GoF on the two-dimensional (2D) areas covered by the optimal and observed regions. The Pareto-optimal regions, originally calculated as sets of discrete points in morphospace, could also be considered as grids of 2D pixels by assuming each optimal point was the center of a small optimal rectangle, facilitating area calculations. Observed species data points were converted to a polygonal region as described earlier (*Survey of observed shapes*). The overlap (or lack thereof) of these two areas was then quantified using Hargrove et al.’s (58) simple goodness of fit parameter:

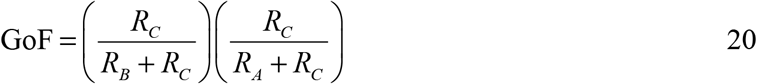

where *R_A_* is the area within the observations but outside the optimal region, *R_B_* is the area within the optimal region but outside the observations, and *R_C_* is the area shared by both the observations and optimal region (Fig. S 15). GoF ranges from 0 for no overlap to 1 for two identical regions, with both lack of overlapped area and extra non-overlapping area being accounted for.

Most of the 148 Pareto-optimal regions we generated yielded very poor GoF, indicating that the corresponding task sets are not constraining the evolution of curved bacterial shapes. The results corresponding to the best six task sets are shown in Fig. S 15. The best two task sets in panels A and B yield nearly identical results but the inclusion of area-weighting with 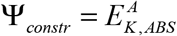 (panel A) results in a cost to elongation of straight rods and thus the logical inclusion of spheres in the optimal region. While the inclusion of 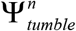 in the task set (panel C, also D and E) yields an acceptable fit, the absence of observed low *ℒ*, high *𝒦* rods predicted in these cases leads us to reject them in favor of the task set corresponding to panel A (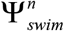, 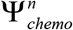, 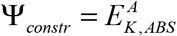).

**Fig. S 1.**
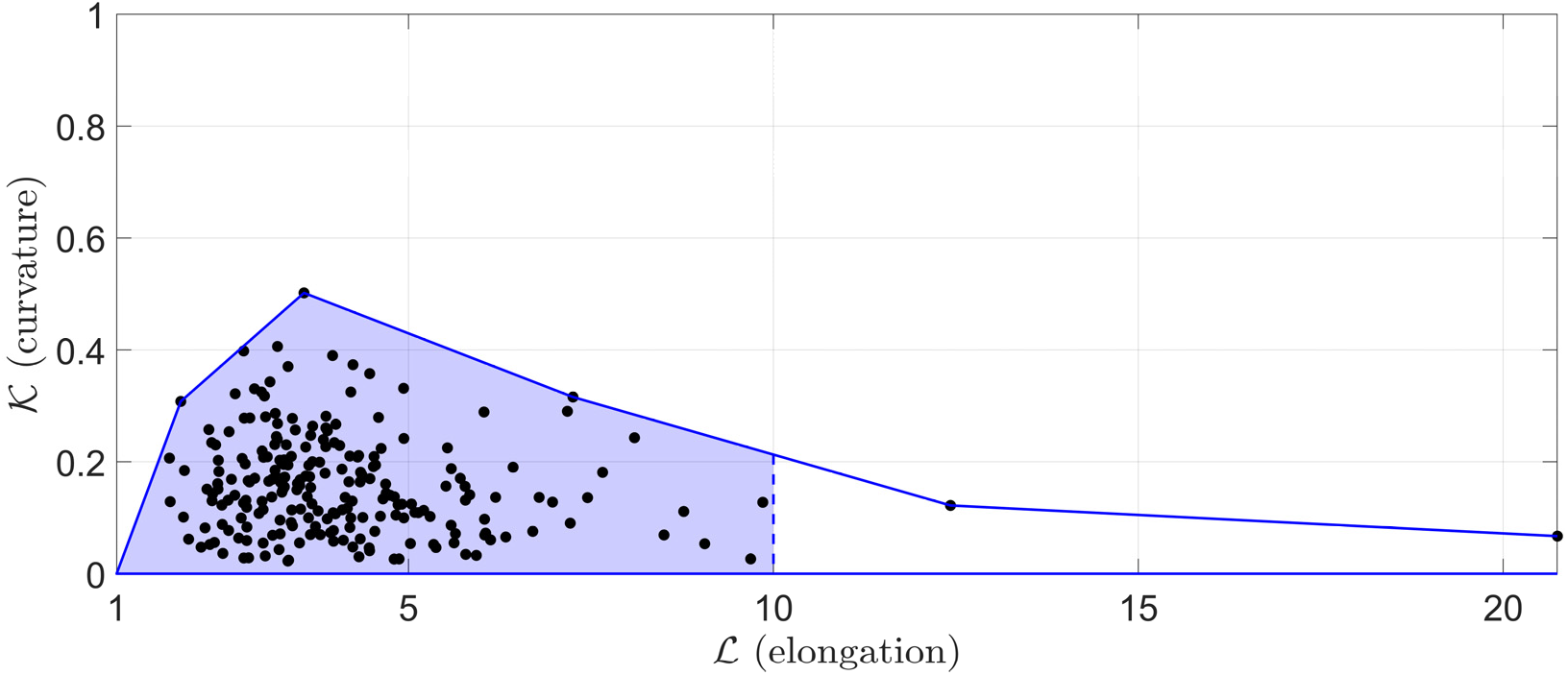
Survey of all observed morphologies (species medians, black dots). Boundary enclosing all points (solid blue line) was computed as described in text; for purposes of computing GoF of the Pareto-optimal region to the observations, this boundary was cut off at the limit of the simulation data, *ℒ* = 10 (dashed blue line), to create a two-dimensional region of observed shapes (shaded blue area).

**Fig. S 2.**
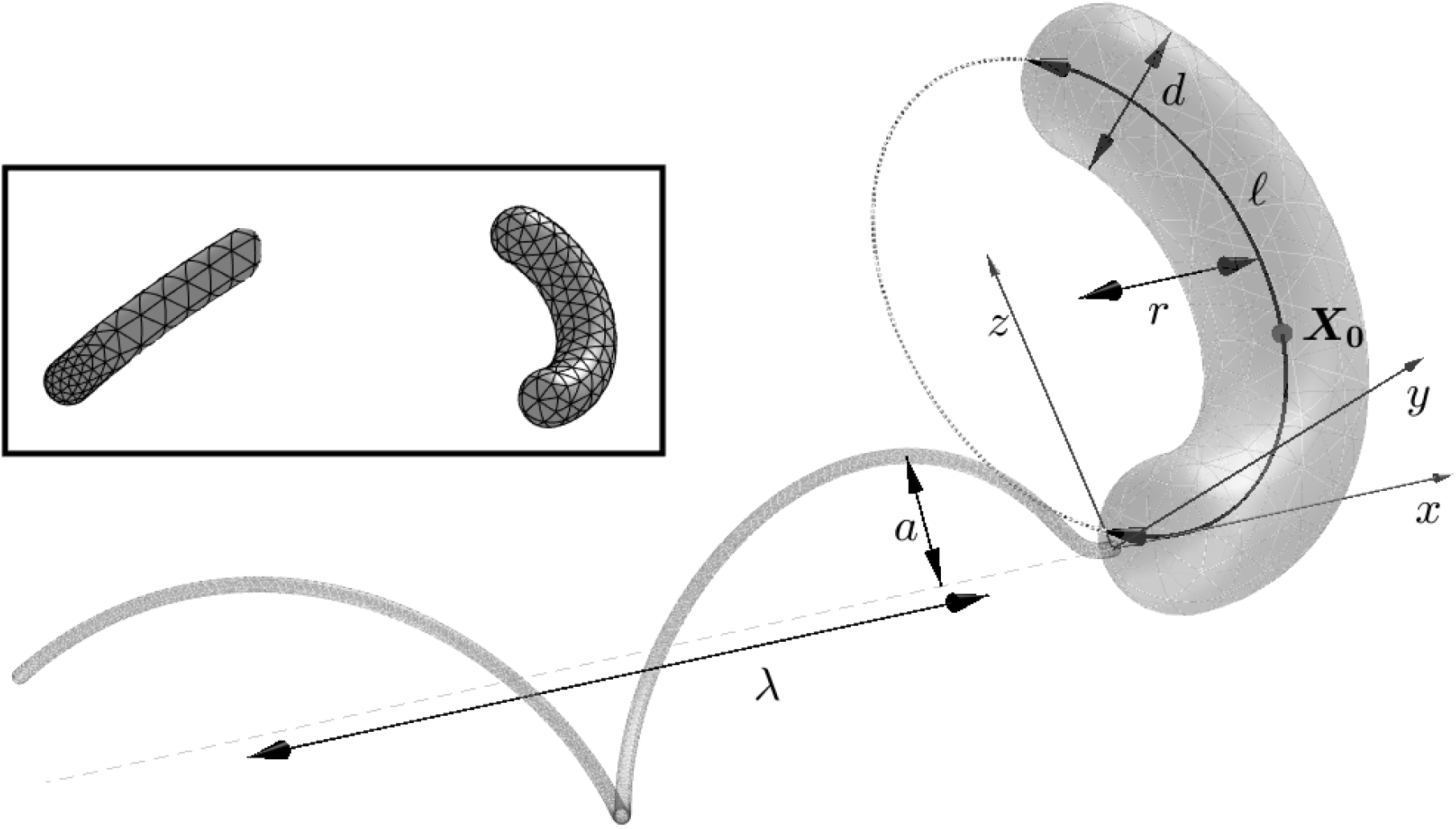
Selected geometric parameters of model curved bacterium (*ℒ* = 5, *𝒦* = 0.5 shown) and flagellum, in the body reference frame. The body centerline is in the x-z plane at t = 0 for free-swimming simulations. Inset: example 2_nd_ order triangular surface meshes of body and distal portion (for clarity) of flagellum.

**Fig. S 3.**
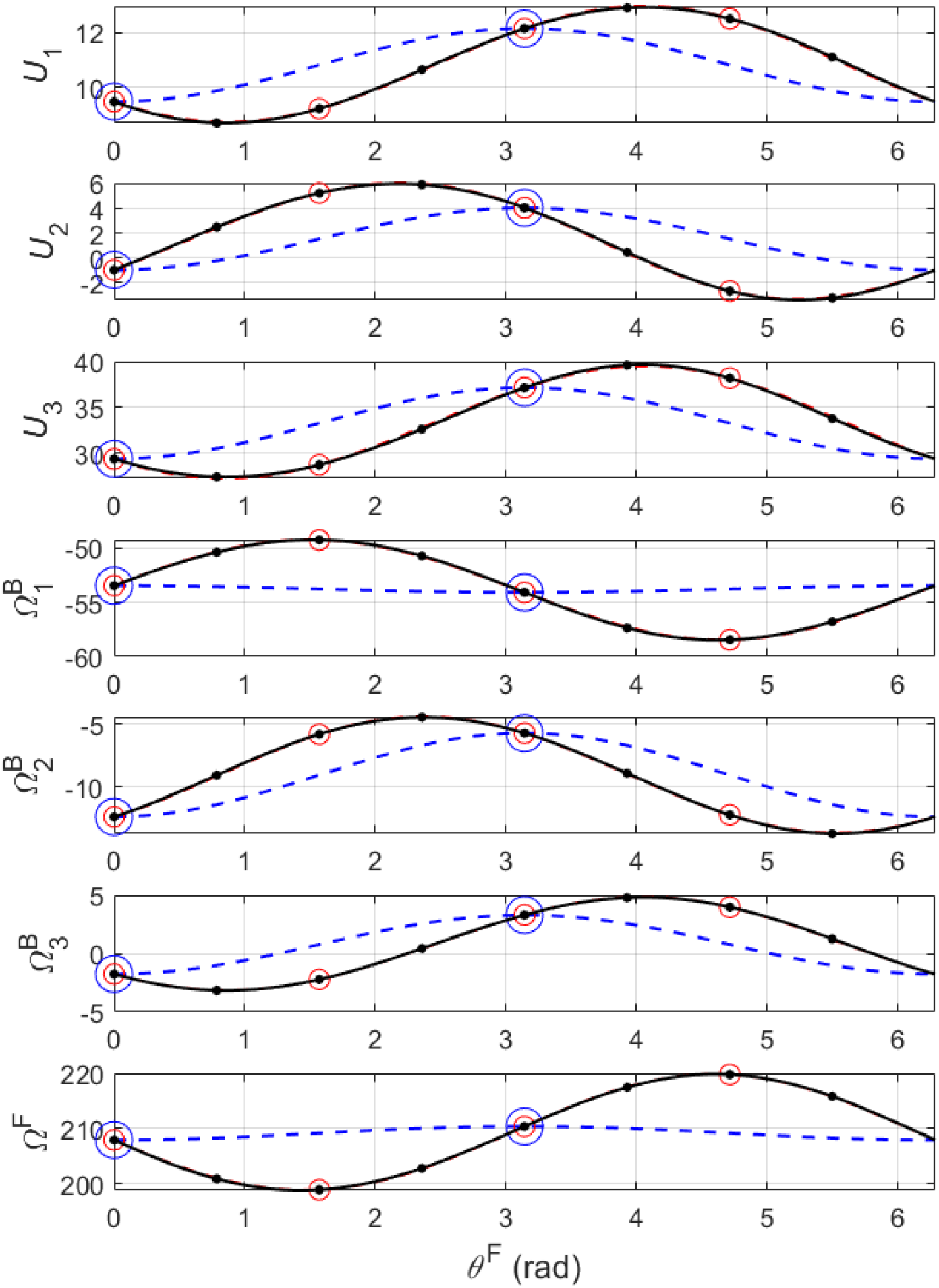
Example Fourier interpolants for the model bacterium shown in Fig. S 2. Interpolants are shown for the kinematic variables ***U*** = [*U*_1_, *U*_2_, *U*_3_] (μm s^-1^), 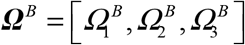 (rad s^-1^), and *Ω^F^* (rad s^-1^) using either two (blue dotted lines and larger circles), four (red dashed lines and smaller circles), or eight (black solid lines and dots) flagellum phase angle (***θ***^*F*^) evaluation points. The interpolants using four and eight points are almost indistinguishable. Since each of these functions must be periodic, their values at, ***θ**^F^* = 2*π*. are immediately known to be the same as those at the first phase angle evaluation *θ^F^* = 0

**Fig. S 4.**
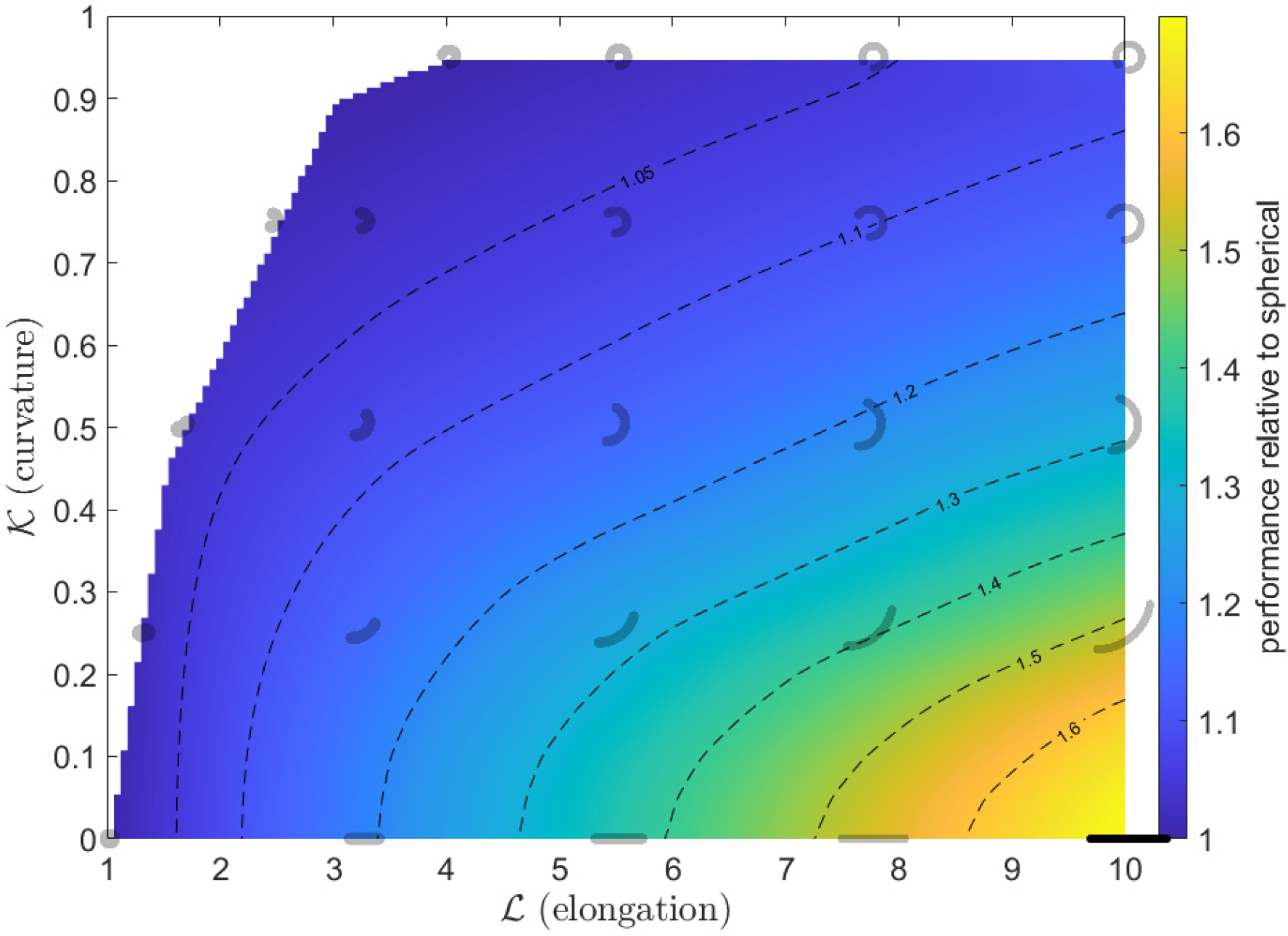
Performance landscape for 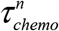, accounting for both body and flagellum and corresponding to flagella optimized for Chemotactic SNR Ψ_*chemo*_. Performance (pseudocolor and contours) is normalized relative to that of a spherical body, i.e., 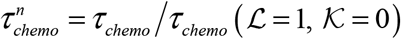. In addition to selected shapes for reference (gray), the performance maximum within our morphospace is shown (black).

**Fig. S 5.**
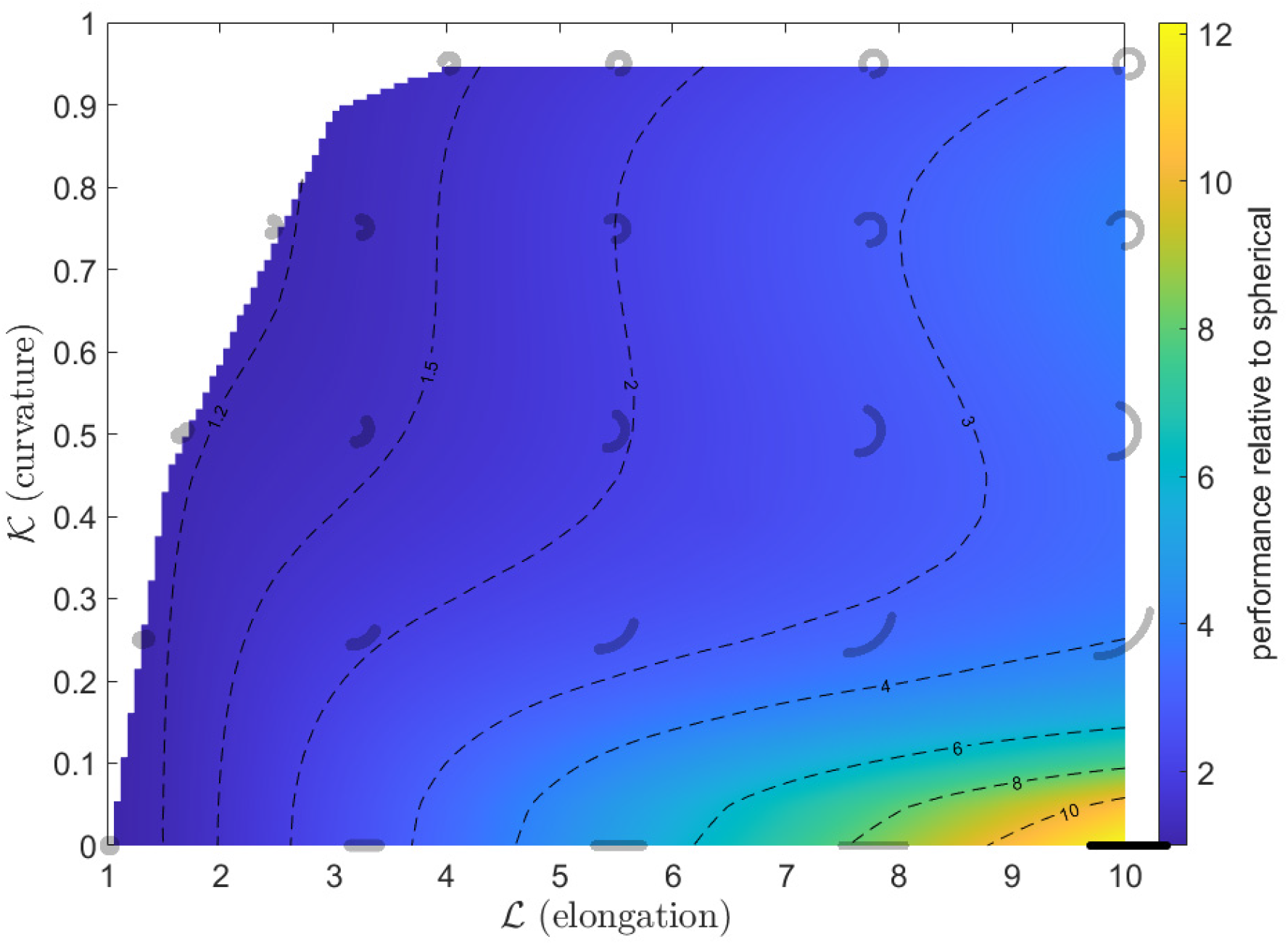
Performance landscape for 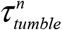, accounting for only the body without flagellum, used for calculation of Tumbling Ease Ψ_*tumble*_. Performance (pseudocolor and contours) is normalized relative to that of a spherical body, i.e., 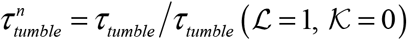. In addition to selected shapes for reference (gray), the performance maximum within our morphospace is shown (black).

**Fig. S 6.**
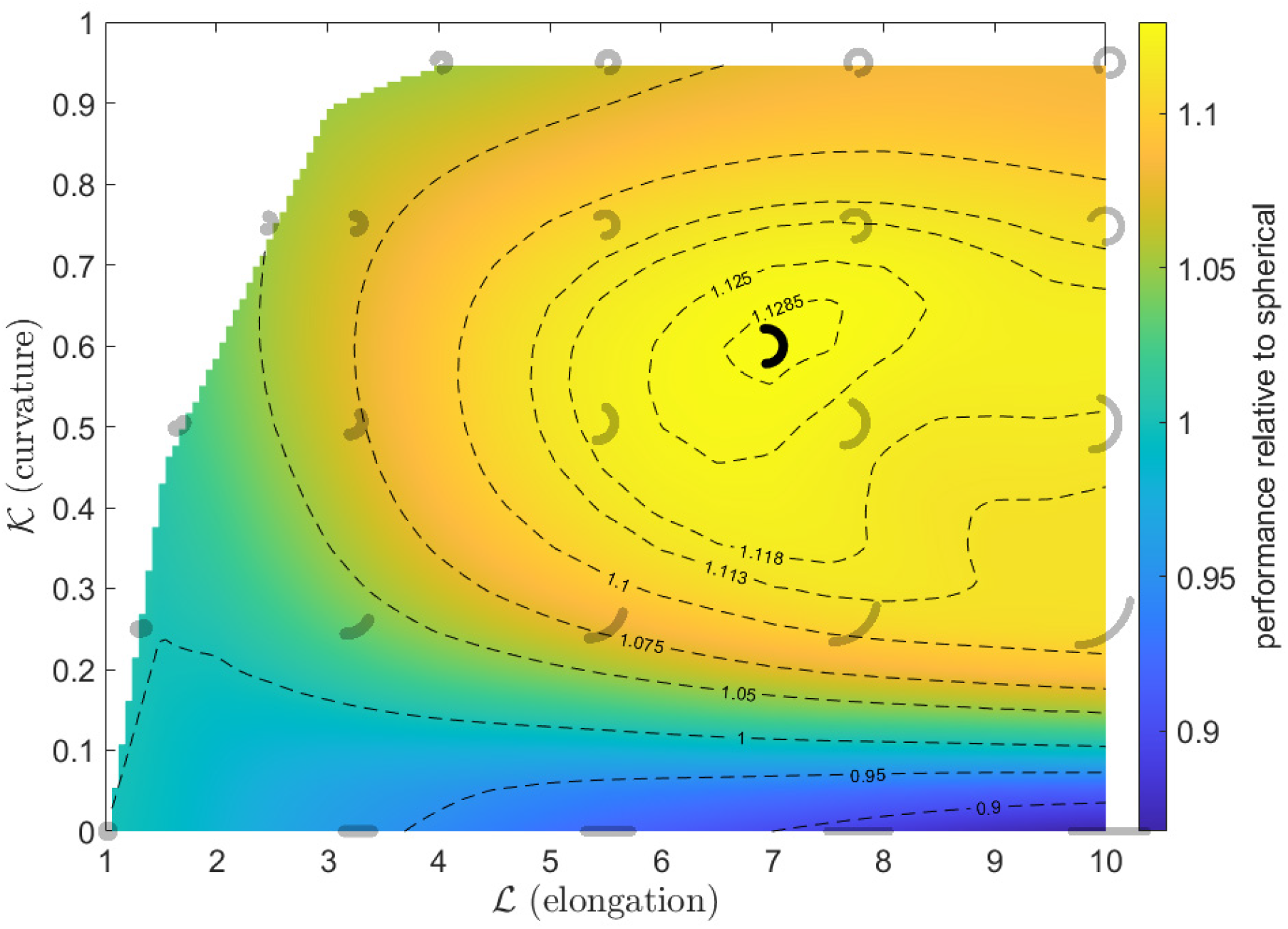
Performance landscape of rescaled effective swimming speed 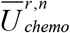 corresponding to flagella optimized for Chemotactic SNR. Performance (pseudocolor and contours) is normalized relative to that of a spherical body, i.e., 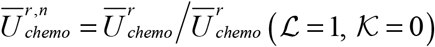. In addition to selected shapes for reference (gray), the performance maximum is shown (black).

**Fig. S 7.**
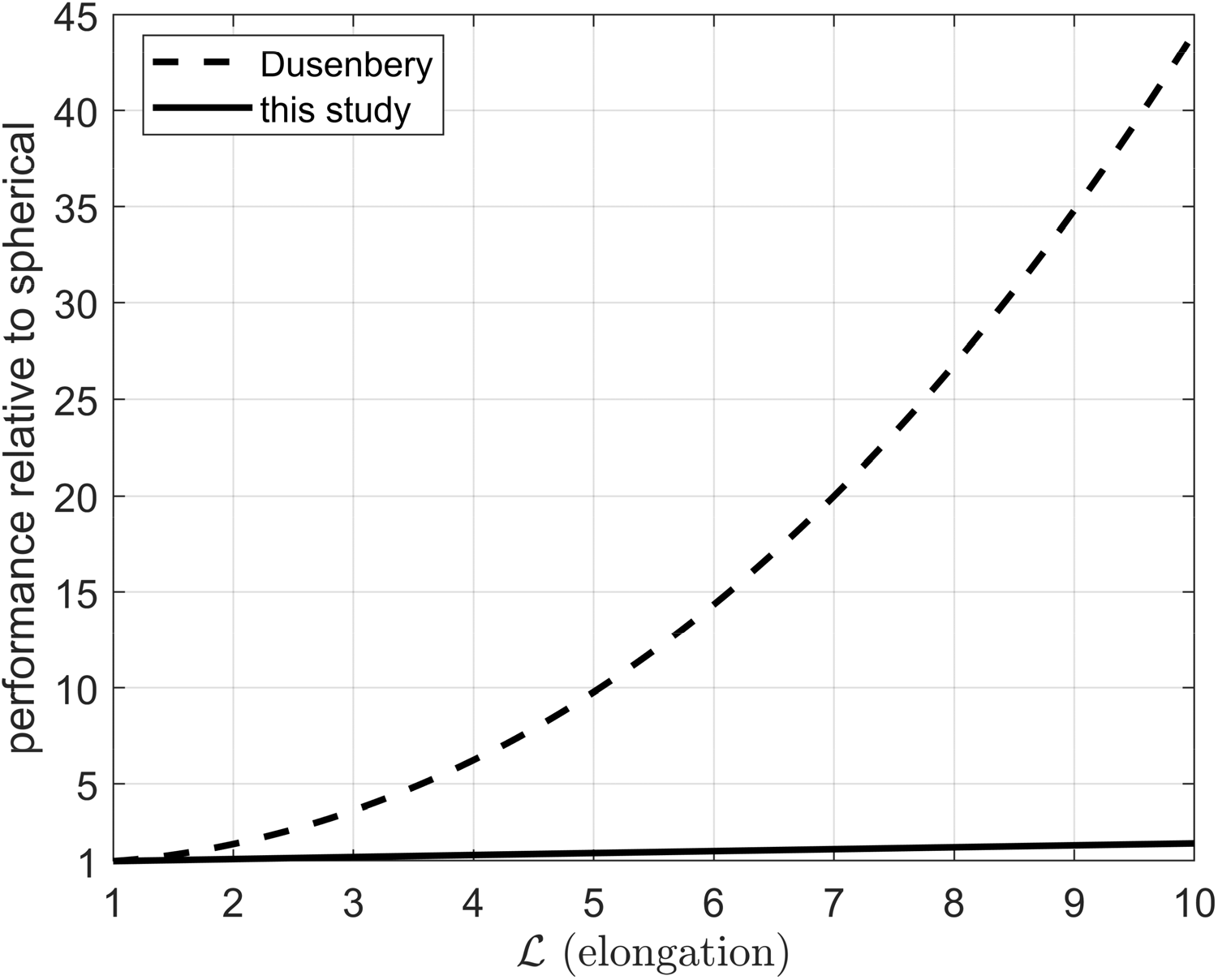
Comparison of normalized Chemotactic SNR 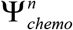 for straight rods (*𝒦* = 0) predicted from this study versus Dusenbery (31). Although both studies predict a monotonic increase with *ℒ*, Dusenbery predicts a factor of 44 improvement at *ℒ* = 10 while we predict a much smaller but still substantial factor of 1.9 improvement. The difference is primarily due to our inclusion of flagellar stabilization against Brownian rotation, which reduces the relative importance of body shape.

**Fig. S 8.**
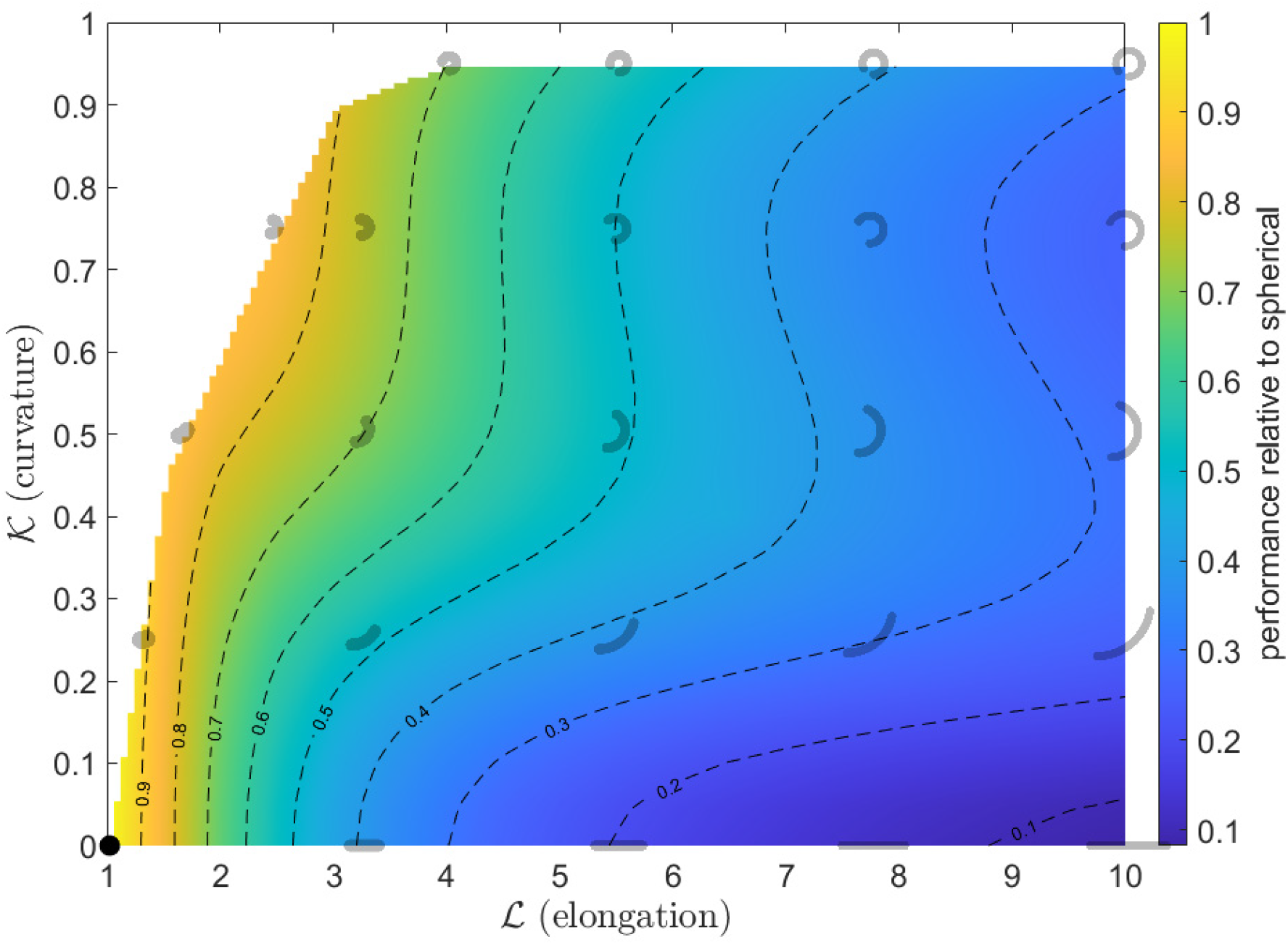
Performance landscape for Tumbling Ease 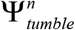. Performance (pseudocolor and contours) is normalized relative to that of a spherical body, i.e., 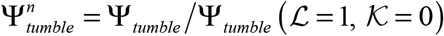. In addition to selected shapes for reference (gray), the performance maximum is shown (black).

**Fig. S 9.**
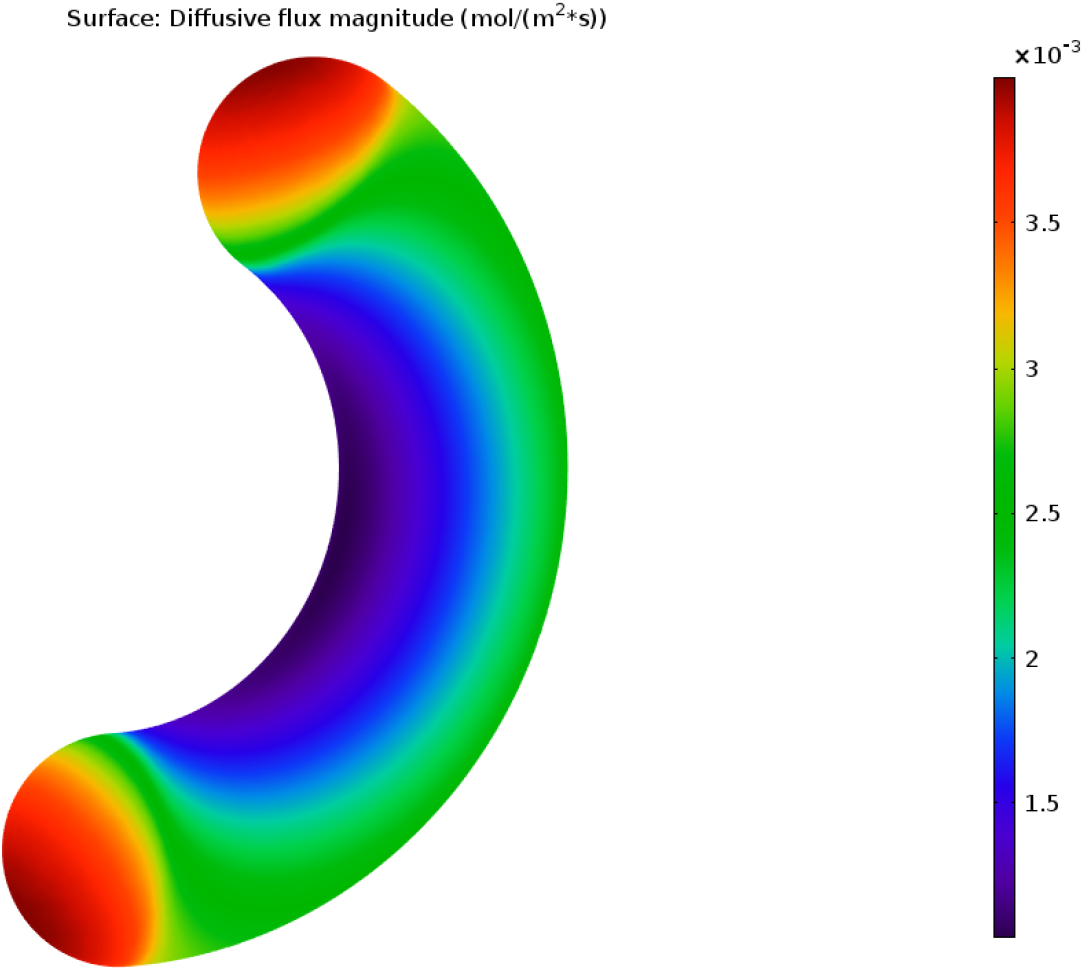
Simulated steady-state diffusive flux of ammonium over the surface of a curved rod, *ℒ* = 5, *𝒦* = 0.5. The scale of flux magnitude here is arbitrary since we later normalize Nutrient Uptake to that of a spherical body.

**Fig. S 10.**
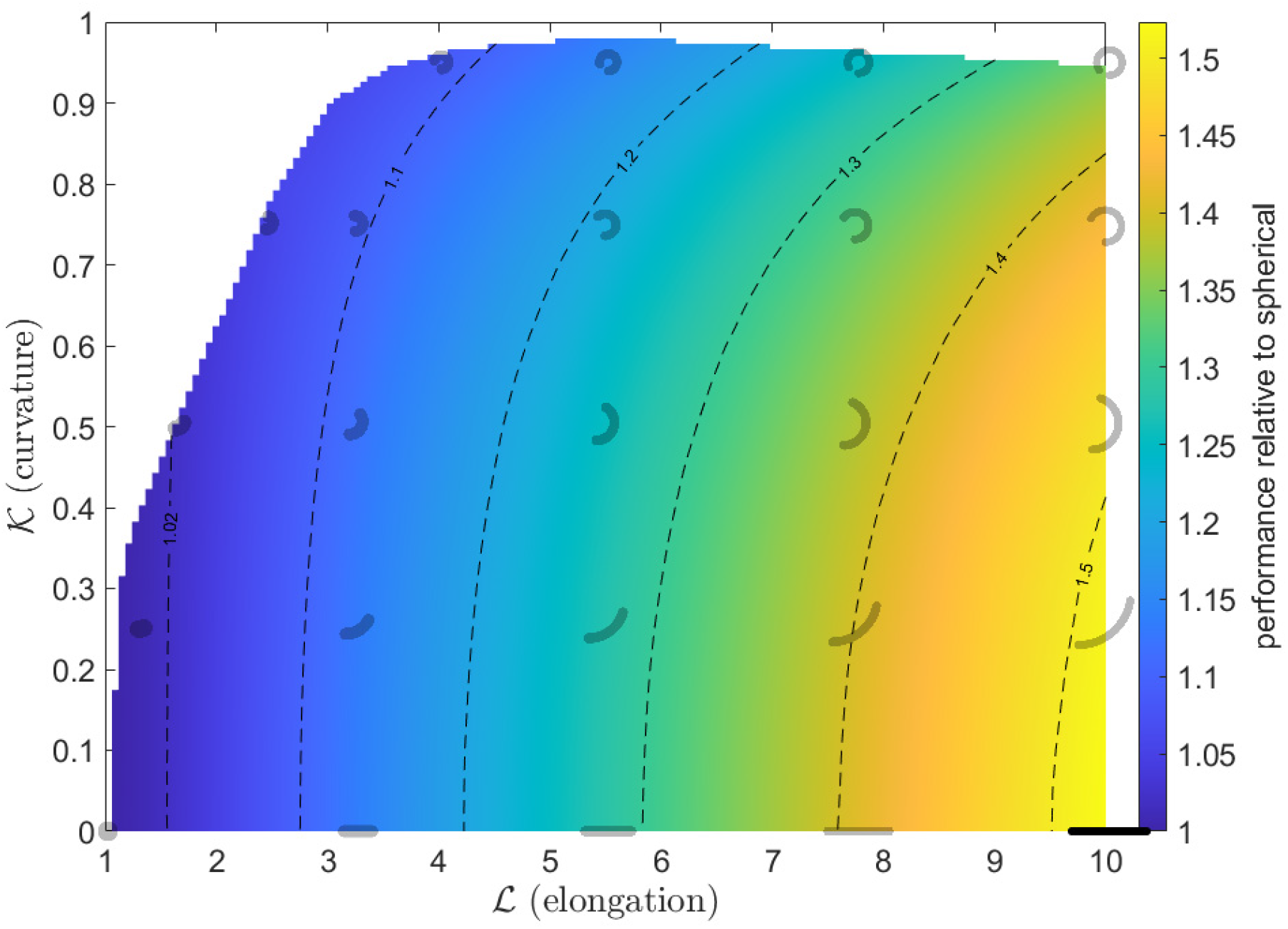
Performance landscape for Nutrient Uptake 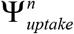. Performance (pseudocolor and contours) is normalized relative to that of a spherical body, i.e., 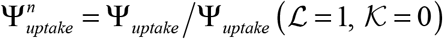. In addition to selected shapes for reference (gray), the performance maximum within our morphospace is shown (black).

**Fig. S 11.**
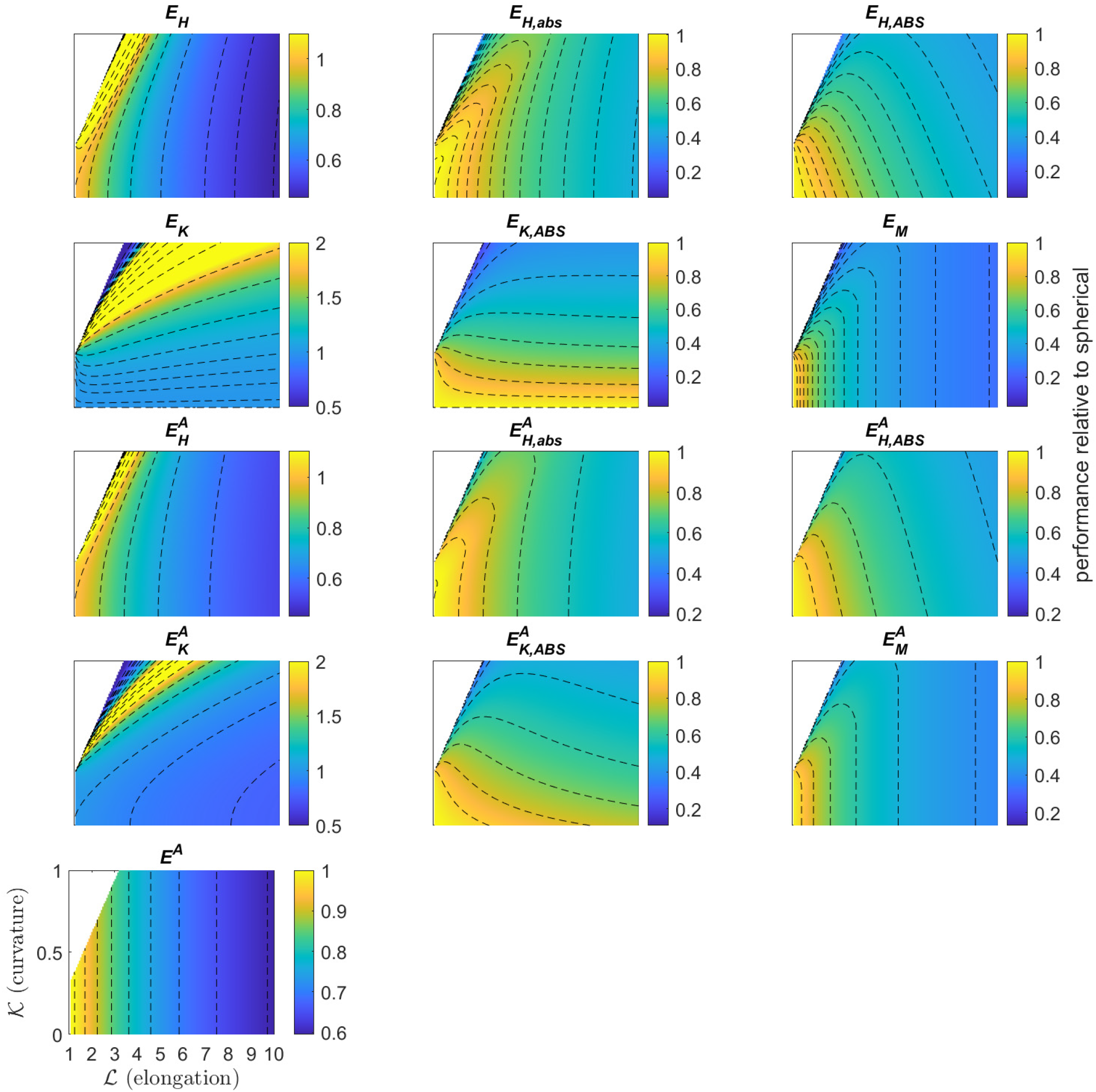
Performance landscapes for putative formulations of Construction Ease 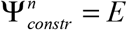. Performance (pseudocolor and contours) is normalized relative to that of a spherical body, i.e., 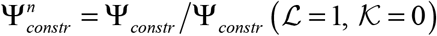. For clarity, contour labels omitted, some colormaps are saturated, and color axes limits vary.

**Fig. S 12.**
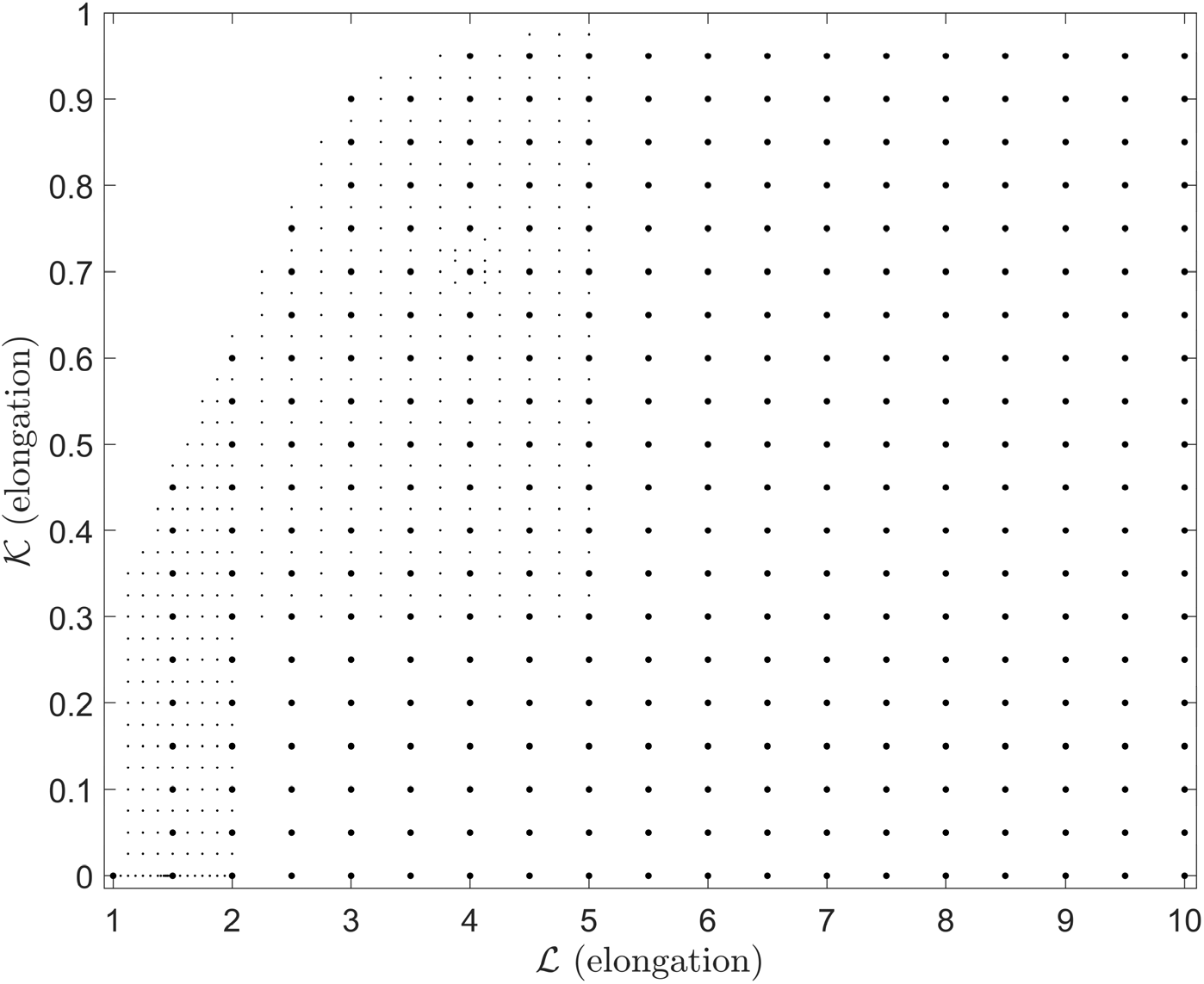
Simulated body shapes used to generate performance landscapes. The baseline grid (larger dots) was used for Chemotactic SNR Ψ_*chemo*_, Tumbling Ease Ψ_*tumble*_, and Nutrient Uptake Ψ_*uptake*_, with additional refined points (smaller dots) used for Swimming Efficiency Ψ_*swim*_. All Construction Ease Ψ_*constr*_ simulations. formulations *E* were defined analytically and did not require discrete

**Fig. S 13.**
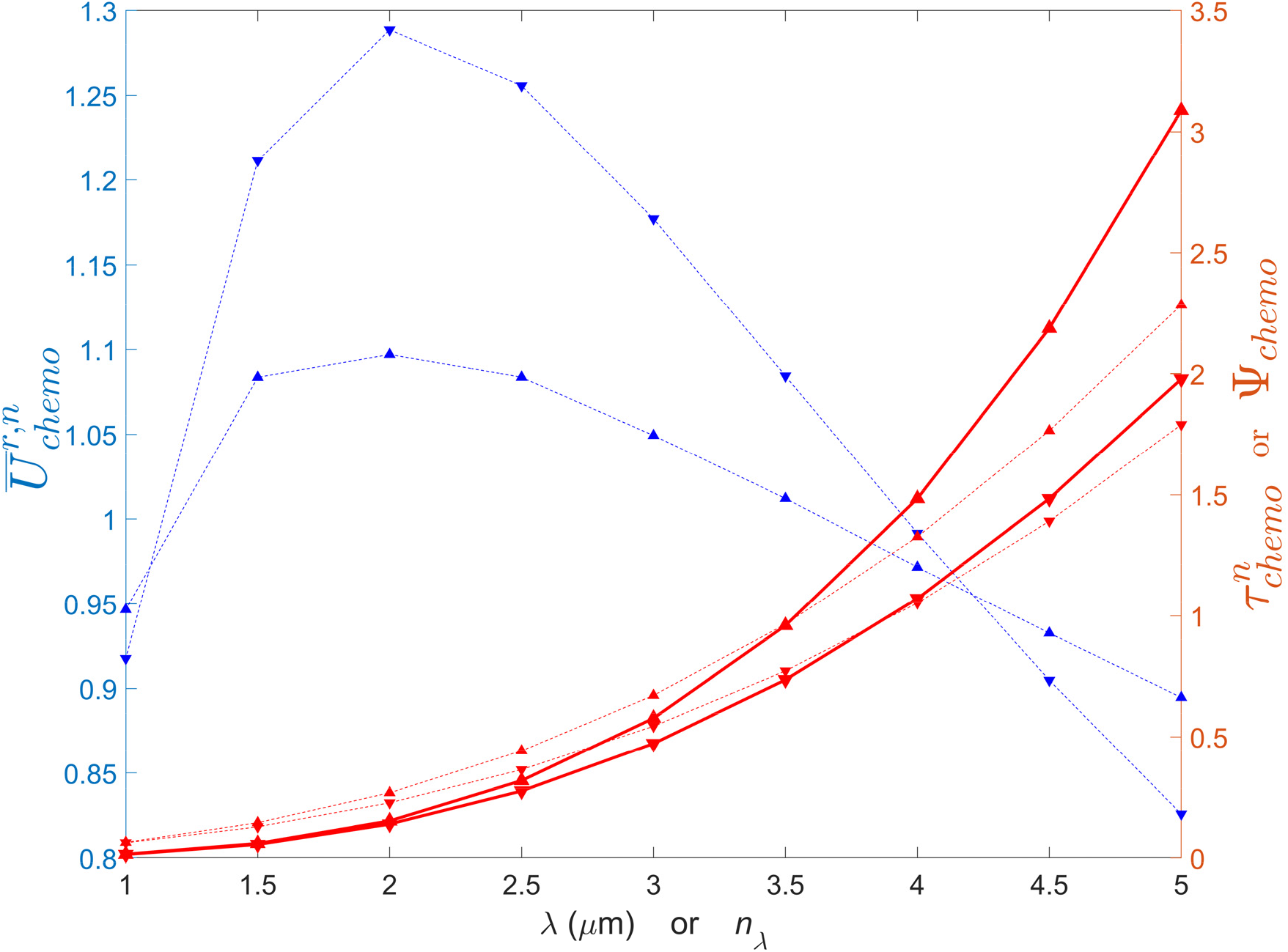
Effect of flagellum length (# wavelengths *n_λ_*, upward triangles and wavelength *λ*, downward triangles) for a spherical body (*ℒ* = 1, *𝒦* = 0) on swimming speed 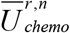 (blue sashed), timescale for loss of orientation 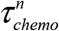 (red dashed), and Chemotactic SNR 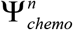 (blue solid). All dependent variables are normalized by their respective values corresponding to the Chemotactic SNR-optimized flagellum for a spherical body shown in Fig. 2C. *λ* was fixed at 3.91 μm while *n_λ_* was varied, *n_λ_* was fixed at 3.55 while *λ* was varied, and flagellum amplitude *a* was fixed at 0.26 μm in all cases - these three values correspond to the flagellum shown in Fig. 2C.

**Fig. S 14.**
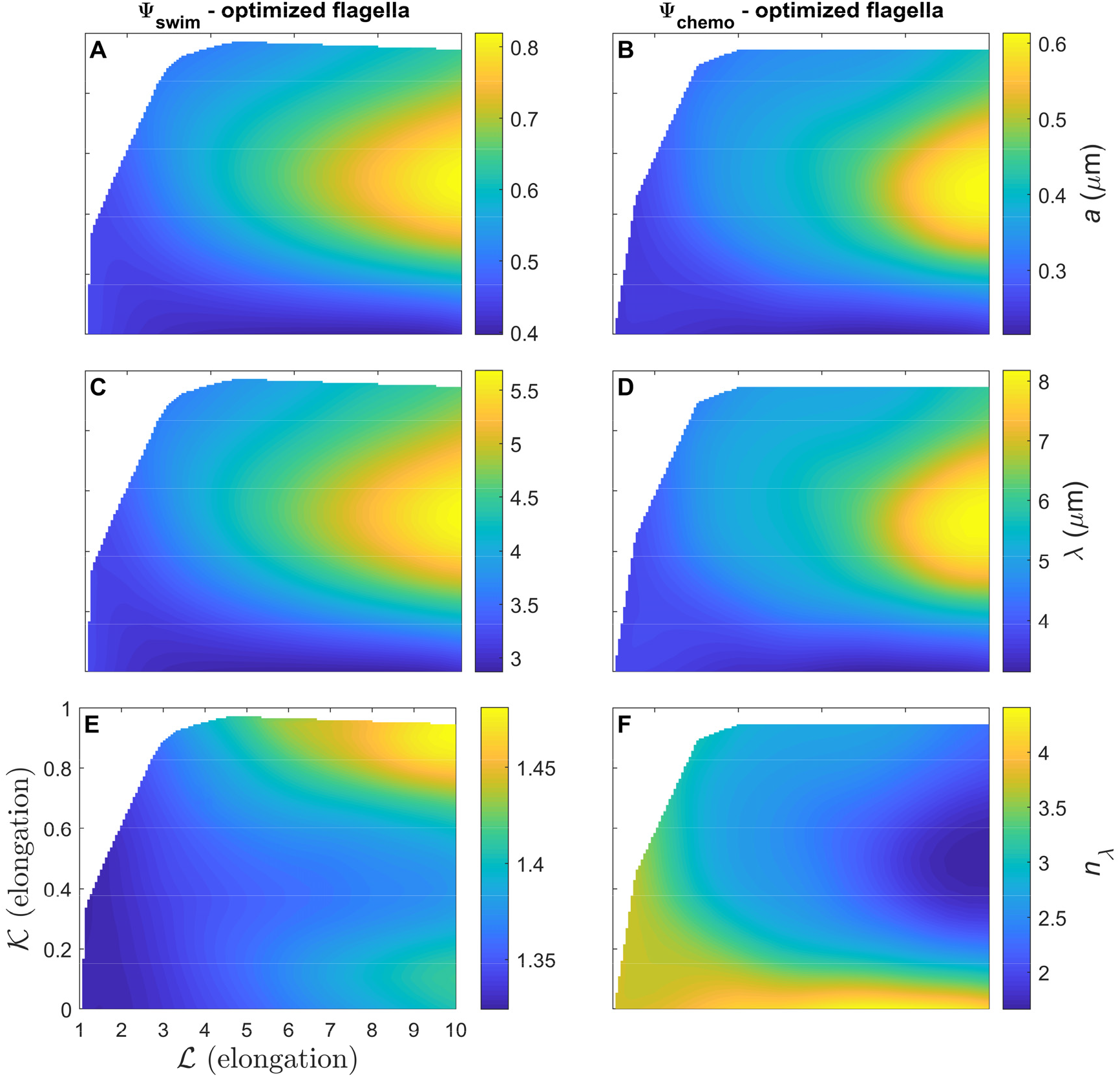
Optimal flagella shape parameters (amplitude *a*, wavelength *λ*, and # wavelengths *n_λ_*) with respect to Swimming Efficiency Ψ_*swim*_ (panels A, C, E) and Chemotactic SNR Ψ_*chemo*_ (panels B, D, F); in the latter case, flagella were constrained to have an arclength of 15 μm. Note varying colormap limits.

**Fig. S 15.**
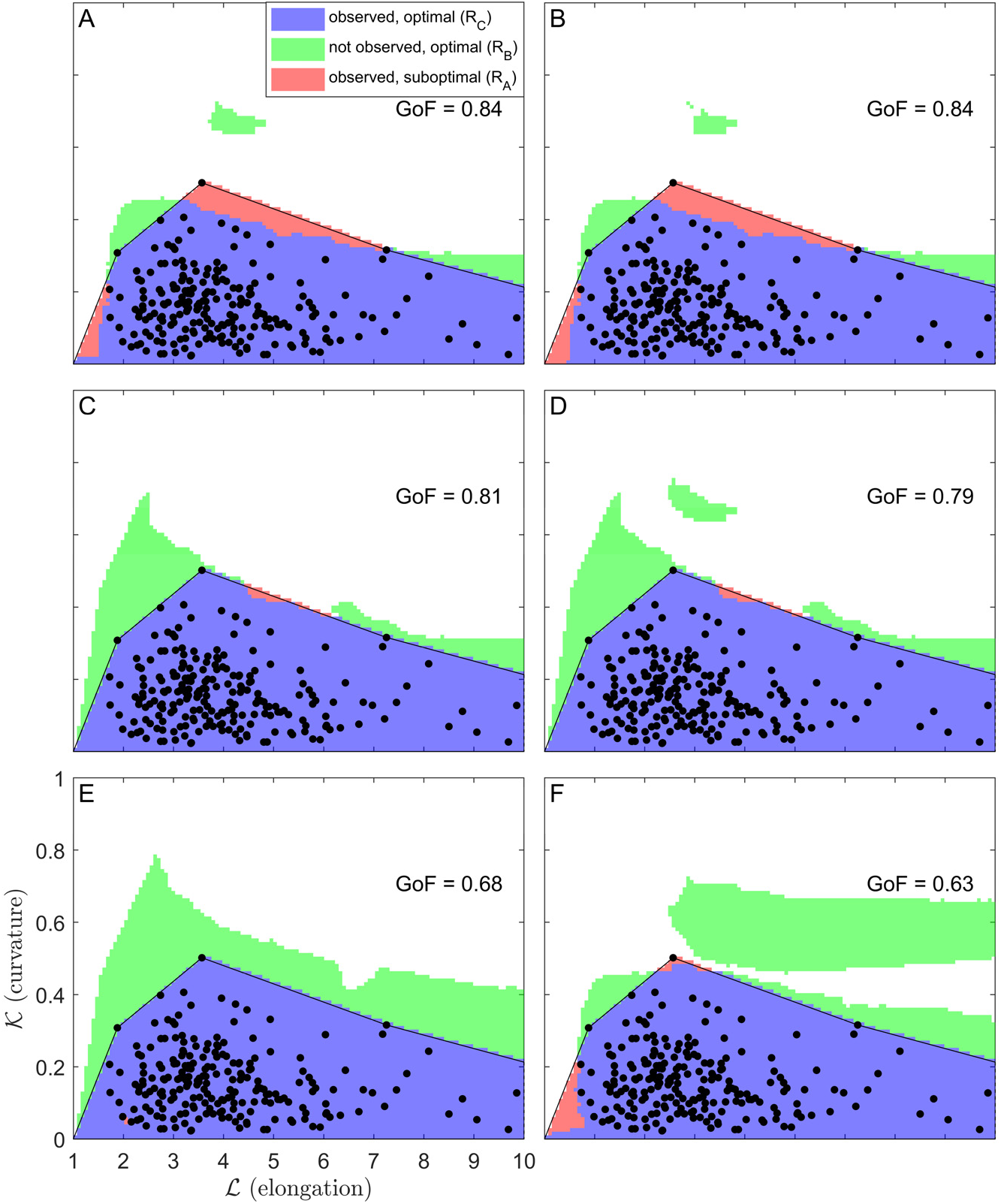
Fits of Pareto-optimal regions resulting from tradeoffs between different sets of tasks to the region of morphospace occupied by observed species medians (dots with black line indicating enclosing boundary). In each case, Goodness of Fit (GoF) was calculated according to equation 20 as a simple function of three areas: the overlapping region that is both observed and optimal, the region that isn’t observed but optimal, and the region that is observed but suboptimal (see SI text). The sets of tasks shown, corresponding to the six highest GoFs of all sets considered, are:

A) 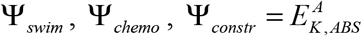 (as in Fig.4, but not smoothed here)
B) Ψ_*swim*_, Ψ_*chemo*_, Ψ_*constr*_ = *E_K,ABS_*
C) 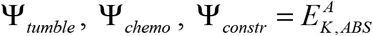 (*E_K,ABS_* yields identical results)
D) 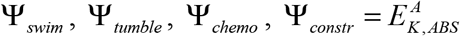 (*E_K,ABS_* yields identical results)
E) 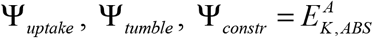 (adding Ψ_*chemo*_ or substituting most other *E* formulations yields nearly identical results)
F) 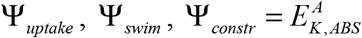 (substituting most other *E* formulations yields nearly identical results)

**Fig. S 16.**
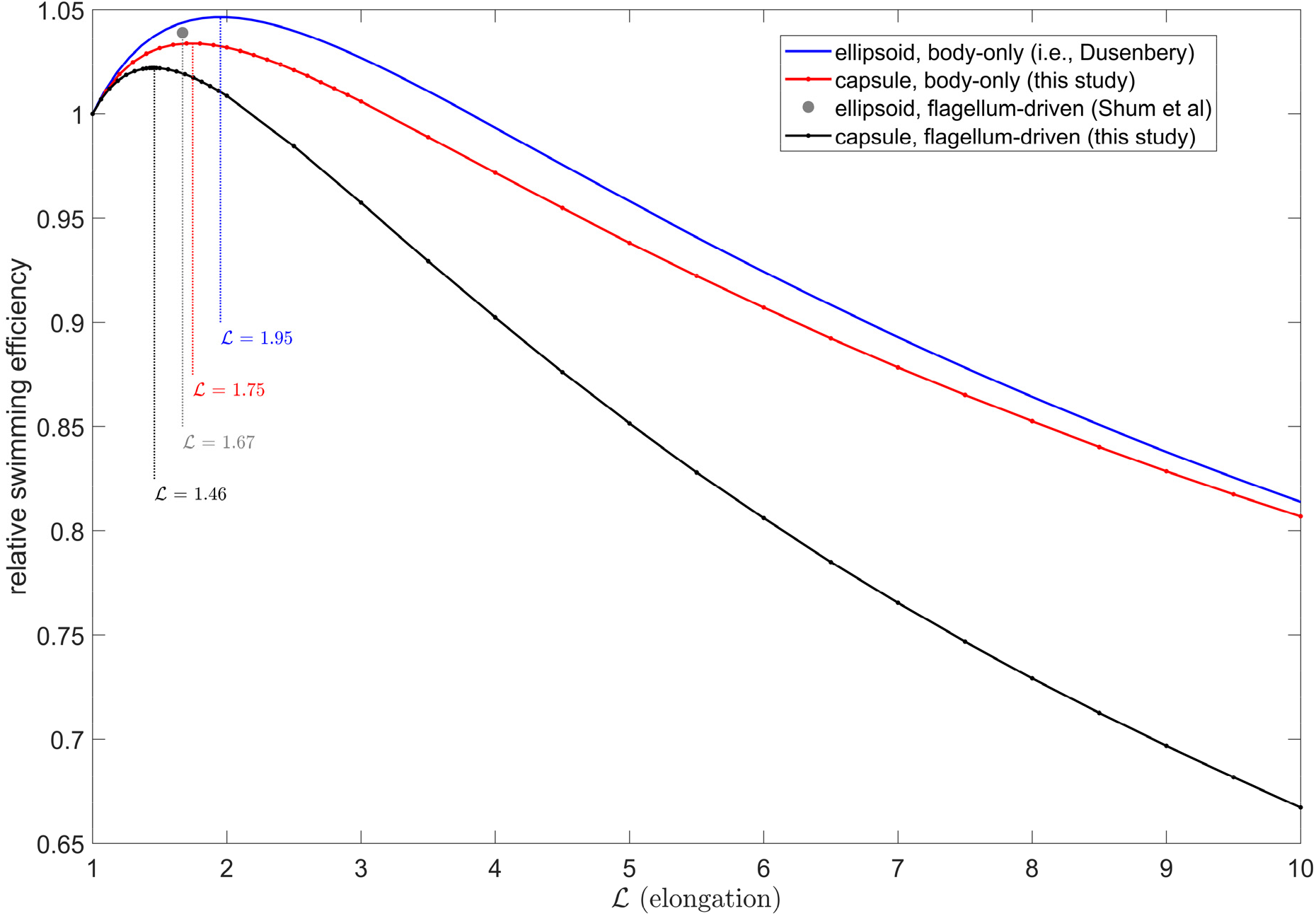
Comparison of swimming efficiency predictions for straight rods modeled as ellipsoids versus capsules, propelled by a rotating flagellum (i.e., equation 14) versus forced without accounting for propulsive mechanism (i.e., assuming efficiency inversely proportional to translational friction coefficient), from this study, Dusenbery (31), and Shum et al. (7). Locations of global maxima are labeled. In all cases, efficiency is normalized to that of a spherical body. Notably, as the biological realism arguably increases within this set of models, the optimal *ℒ* decreases toward spherical.

**Fig. S 17.**
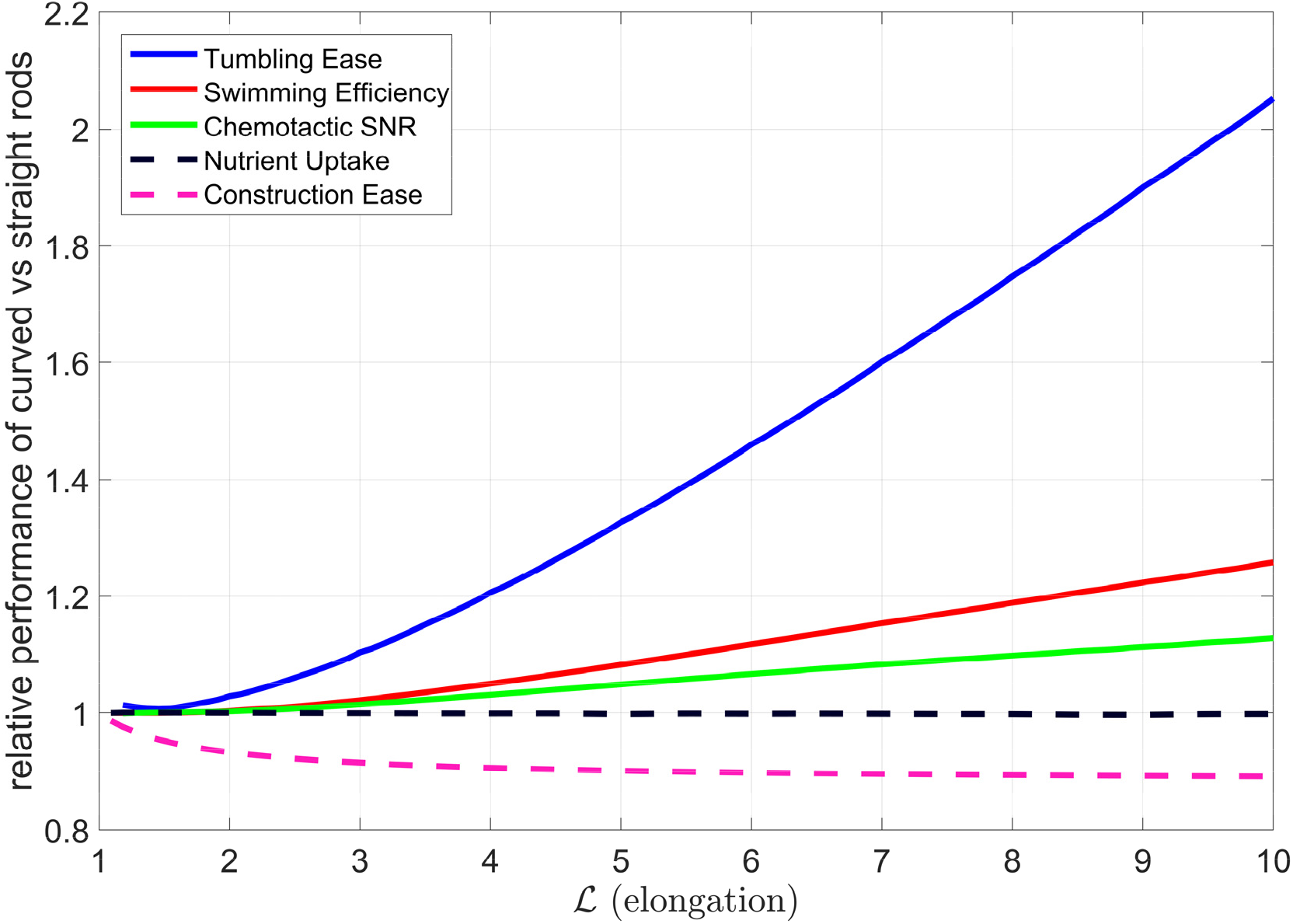
Ratio of performance of curved (*𝒦* = 0.14, i.e. the species median) to straight rods (*𝒦* = 0) at different tasks, calculated by evaluating the interpolants for each task along these transects. Some tasks are improved by curvature (solid lines) while others become worse (dashed lines).

**Table S 1.**
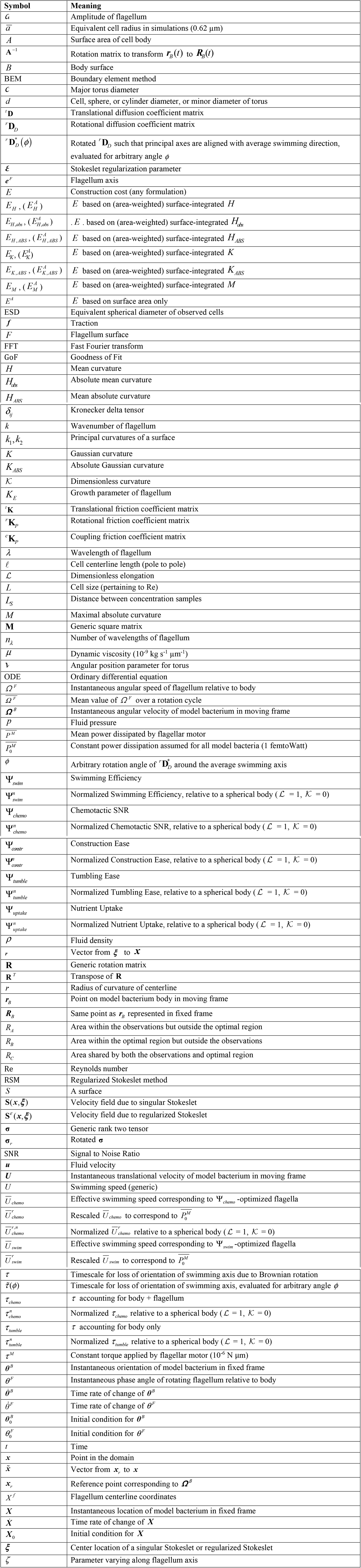
Definitions of symbols and abbreviations used in this paper.

**Table S2.** Curved rod morphological survey data for each image in our dataset, assembled using MicrobeJ (2). The 2_nd_ sheet, *aggregated images*, is a list of the sets of images for each species that were aggregated when taking medians due to having the same strain, culture conditions, etc.

**Table S3.** Curved rod morphological survey data, aggregated to the level of species medians, used to create Fig. 1 and Fig. S 1.

**Movie S1.** “Race” between several model bacteria paired with their efficiency-optimized flagella, their temporal swimming kinematics rescaled such that each dissipates the same flagellar power 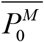, and their trajectories aligned in space. Morphologies are in order of decreasing Swimming Efficiency **ψ**_*swim*_: from top to bottom, (*ℒ*, *𝒦*) = (1.46, 0), (4, 0.7), (1, 0), (5, 0.975), (10, 0.5), (10, 0). Note the lack of correlation between Swimming Efficiency and the geometry (e.g. amplitude) of the swimming trajectory.

